# Latent neural population dynamics underlying normal breathing, opioid induced respiratory depression, and gasping

**DOI:** 10.1101/2022.11.30.518585

**Authors:** Nicholas E Bush, Jan-Marino Ramirez

## Abstract

Breathing is vital and must be concurrently robust and flexible. This rhythmic behavior is generated and maintained within a rostro-caudally aligned set of medullary nuclei called the Ventral Respiratory Column (VRC). The rhythmic properties of individual VRC nuclei are well known, yet technical challenges have limited the interrogation of the entire VRC population simultaneously. Here, we characterize over 13,000 VRC units using high-density electrophysiology, opto-tagging, and histological reconstruction. Population dynamics analysis reveals interactions between inspiratory and expiratory dynamical states give rise to a low-dimensional, elliptical neural manifold. The structure of this manifold is robust and maintained even during opioid-induced respiratory depression (OIRD). However, during severe hypoxia-induced gasping, the low-dimensional dynamics of the VRC reconfigure from rotational to all-or-none, ballistic efforts.

## Main

Recent advances in the ability to record the activity of large populations of neurons has driven concurrent adoption of dimensionality reduction techniques to describe complex population dynamics^1–4^. These approaches have been widely successful across many systems to uncover simple attractor-like dynamics that describe neural population space.

Additionally, studies from small invertebrate neuronal networks have demonstrated that qualitatively and quantitatively similar patterns of rhythmic activity can be observed over a large regime of varying combination of biophysical properties, suggesting that neuronal networks maintain specific patterns of activity that can emerge from many different sets of synaptic strengths and intrinsic membrane properties^5^. That is, numerous solutions for a given neural behavior can arise from a degenerate space of constituent cellular components.

The neural circuits that drive and maintain breathing in mammals afford a tractable handle on the intersection of these two complimentary descriptions of redundancy underlying neural computations. Breathing is a rhythmic, stereotyped motor behavior that is constitutively maintained throughout the life of an animal. Despite the relative simplicity of this behavior, the neural circuits that drive and maintain breathing are distributed throughout a large population of anatomically distributed and molecularly diverse neural population that compose the rostro-caudally extended Ventral Respiratory Column (VRC)^6–10^. The VRC contains vital centers that act as the scaffold for maintaining breathing, including the intrinsically rhythmic, bilaterally synchronized, preBötzinger Complex (preBötC)^11–14^, the chemo-sensitive retrotrapezoid nucleus (RTN, also termed the parafacial respiratory group, pFRG)^15^, and bulbospinal premotor neurons of the ventral respiratory group (VRG)^16^. The VRC also participates in critical bidirectional integration with medullary, pontine, thalamic, and cortical centers^17–19^, as well as the peripheral nervous system^20,21^. This vast interconnectivity imbues the respiratory centers with sufficient flexibility to support varied control of breathing during behaviors such as whisking and vocalizing^22,23^.

Here, we introduce a novel experimental preparation that allows for large-scale electrophysiological recordings along the rostrocaudal extent of the medulla. We combine recordings from Neuropixel probes with optogenetic tagging^24,25^ and 3D histological reconstructions^26^ to detail the respiratory-related activities of VRC neural populations in intact, freely breathing mice. We test the hypothesis that VRC-wide neural populations exhibit simple, rhythmic and stereotyped low-dimensional dynamics. Our results show that VRC population activity evolves along a continuous, rotational, low-dimensional trajectory that is consistent across animals and recordings. Inspiratory and expiratory activity alternates between disjoint, but intersecting, linear dynamical systems governed by unstable fixed points. Notably, the offset of inspiration forms an attractive target in neural population space, thus expanding the classical idea of the “inspiratory off-switch”^27–29^ to the population dynamics of the entire VRC.

We further test how these rotational dynamics are disrupted by systemic, physiologically relevant perturbations to respiratory dynamics: opioids and hypoxia. Opioid-induced respiratory depression (OIRD) is a leading cause of death in the United States^30,31^ and our understanding of the molecular, cellular, and circuit mechanisms of OIRD is continually evolving^32–35^. Here we show that opioids cause myriad diverse changes in single unit spiking activity across the VRC. However, the underlying low-dimensional structure is preserved, suggesting that there are redundant network solutions to preserve respiratory function in response to perturbations.

In contrast, acute hypoxia induces dramatic physiological pressure to alter the normal breathing pattern to produce auto-resuscitative gasping^36,37^. The failure to rouse during gasping underlies susceptibility to Sudden Unexpected Infant Death syndrome (SUIDS)^38–40^. During gasping, the rotational dynamics observed during eupnea (normal breathing) collapse to form ballistic all-or-nothing efforts.

Together, these data reframe the neural circuits that govern breathing in the context of the emerging field of population dynamics. VRC populations can be described by simple low-dimensional dynamics that are consistent across animals.

## Results

### High resolution mapping of anatomically and optogenetically identified neurons in the VRC

Neuropixel probes^41^ were inserted along the rostrocaudal axis of the VRC of 30 urethane anesthetized mice to record single unit activity from the entire VRC simultaneously (Fig. 1a, Extended Data Fig. 1). Urethane anesthesia sustains respiratory rates close to those observed in awake animals (approximately 2-4 Hz) and maintains many cardiorespiratory reflexes. Breathing was monitored with diaphragm electromyogram (EMG). Across all animals, we recorded 13,336 single units in 116 separate recordings (average 3.8 recordings per animal, 115 units per recording). Repeated insertion of the Neuropixel probe did not severely alter respiratory activity (Extended Data Fig. 1). We identify units as putatively axonal or somatic in origin based on their extracellular waveform shape as recently demonstrated by Sibille et al.^42^ (Extended Data Fig. 2). Fluorescent DiI was used to identify the locations of each probe insertion in post-hoc histological analyses (Fig. 1b, Extended Data Fig. 1). In addition, we use opto-tagging^24^ to identify a subset of recorded neurons by transcriptional markers (Extended Data Fig. 2h-j). We recorded 329/2504 (13.1%) Vglut2^+^; 251/2866 (8.8%) ChAT^+^; 554/5774(9.6%) Vgat^+^; and 542/2192 (24.7%) Dbx1^+^ units. Dbx1^+^ units are of interest as they are critically involved in maintaining breathing across the VRC^11,16,43^.

**Figure 1.**
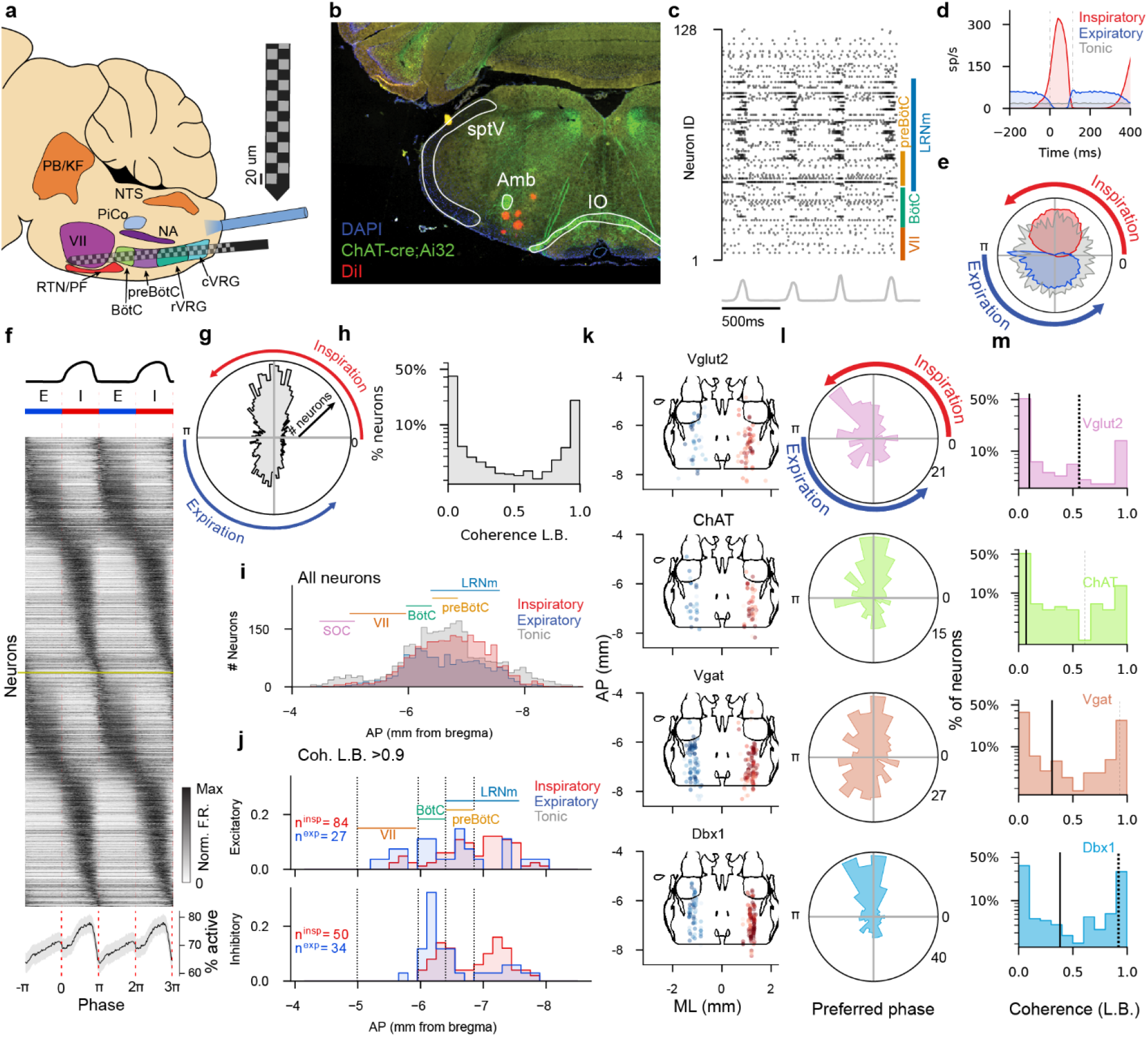
Large scale recording to characterize VRC populations. (**a**) Schematic of recording technique. Neuropixel probe (detail in inset, gray boxes are active sites) is inserted into the VRC of the left medulla of urethane anesthetized mice. All data are for units identified as somatic (Extended data Fig. 2). (**b**) Example coronal section showing cross sections of recording tracks (red) from 5 sequential VRC recordings. Colored bars show anterior-posterior extent of the indicated anatomical markers. (**c**) (top) Raster of 128 units with corresponding integrated diaphragm muscle EMG (gray) over 1.5 seconds. (**d**) Diaphragm onset aligned peri-event time histogram (PETH) for 3 units (red: inspiratory, blue: expiratory, gray: tonic). Mean F.R. +/− S.E.M. is shown, but S.E.M. is small enough to not be visible. (**e**) Phasic tuning curves of the 3 example units in (d). Radial axis is spike rate normalized to maximal spike rate, angular axis is phase of breath; Breath onset is 0, breath offset is *π*. (**f**) Phase aligned, maximum normalized responses of all phasic, somatic units (Coherence L.B.>0.1) recorded across all recordings (n=3,637). Responses are duplicated in both axis directions to visualize phasic boundaries. Bottom shows percent of neurons active at a given point in the breathing cycle, where active is defined as >10% max firing rate. (**g**) Angular histogram of preferred phase of all phasic units. (**h**) distribution of the lower bound of coherence for all units recorded (n=6,315) (**i**) Marginal distributions of all inspiratory (red), expiratory (blue) and tonic (gray) units recorded along the anterior-posterior (AP) axis (binsize is 100μm). AP boundaries of some nuclei in the ventral medulla are shown (SOC-Superior Olivary Complex; VII: Facial nucleus; BötC – Bötzinger Complex; preBötC – pre-Bötzinger Complex; LRNm – Lateral reticular nucleus, medial division) (**j**) AP distributions of strongly coherent units (*C_lb_* > 0.9) identified by optotagging as excitatory (either Vglut2^+^ or Dbx1^+^, top) or inhibitory (Vgat^+^, bottom) Red is inspiratory units, blue is expiratory units. (**k**) Location and respiratory coherence of all positively tagged, phasic units. Expiratory units are shown on the left, inspiratory units on the right for visualization; all units were recorded from the left medulla. Dot saturation indicates the coherence of that unit (darker units are more coherent). (**l**) Angular histograms of preferred phase for all phasic, positively tagged units. Angular axis is preferred phase lag (i.e., when in a breath the unit is active), and radial axis is number of units with a preferred phase lag. (**m**) Distributions of lower bound of coherence for all positively tagged units. Black vertical line is median, dashed vertical line is 75^th^ percentile.

**Figure 2.**
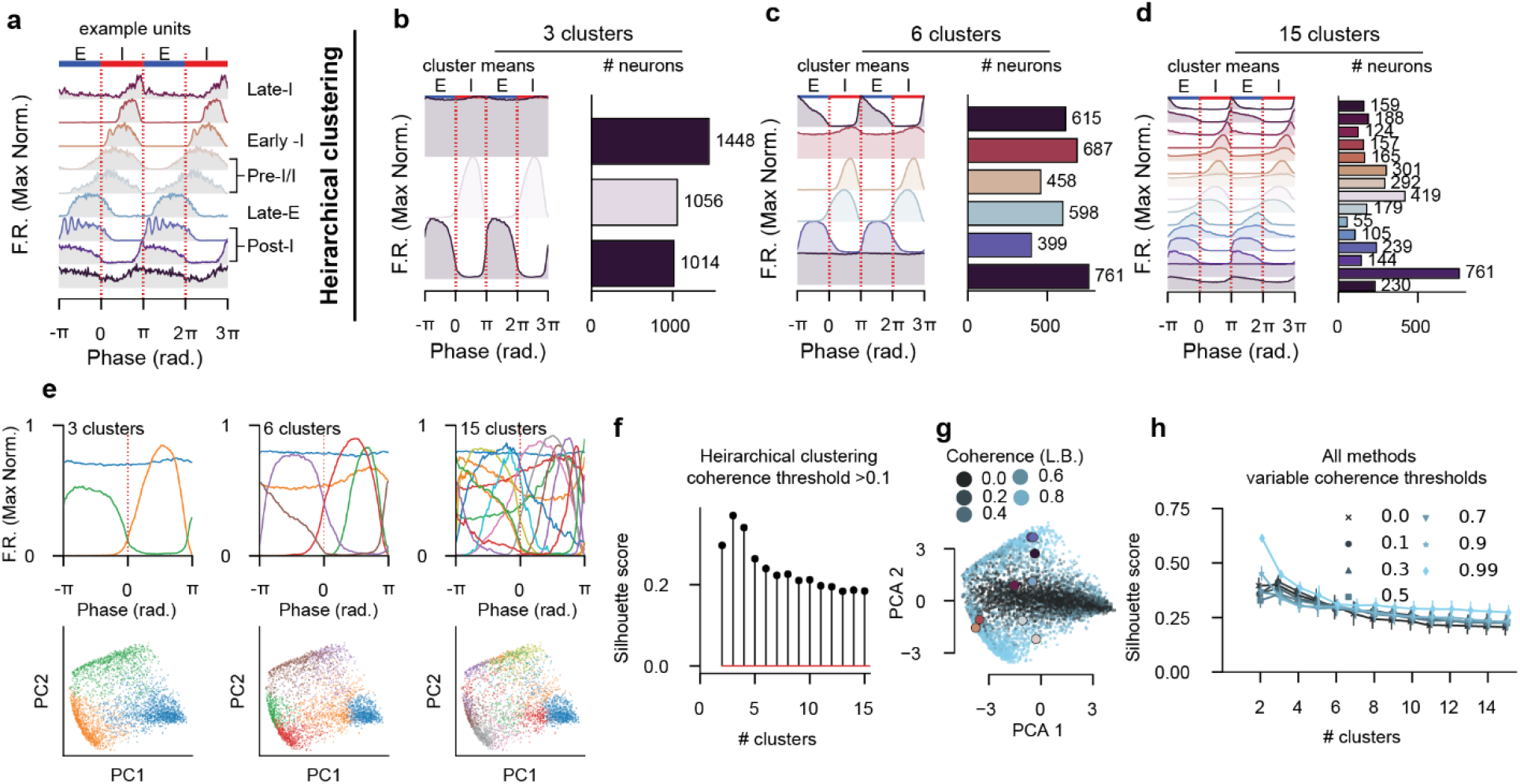
Clustering of phasic units reveals continuum of activity patterns. All data are for somatic units only. (**a**) Nine example phasic (coherence lower bound >0.1) units with classical activity patterns with respect to breathing phase. Firing rates are maximum normalized. Classical activity pattern identities are given right based on manual identification. (**b-d**) Hierarchical clustering of phasic activity patterns showing (left) the mean activity pattern for a given cluster, and (right) the number of units assigned to that cluster for clustering into 3,6, and 15 clusters. Separating units into three clusters results inspiratory, expiratory, and tonic neurons; six clusters results in clusters that resemble classical post-inspiratory, pre-inspiratory, inspiratory, expiratory, and tonic units, while 15 units results in a continuum of clusters that tile the breathing phase with both incrementing and decrementing patterns^6,7,27,46^ with varying phasic structure. (**e**) (top) Cluster means and (bottom) projection of phasic tuning curves into two-dimensional PC space, colored by cluster identity as in (top). Each dot is a unit. Clustering for 3, 6, and 15 clusters are shown. (N.B.: clustering here is performed on the raw phase average responses, and the PC projections are for visualization). (**f**) Maximal silhouette score for hierarchical clustering for 2 through 15 computed clusters. The maximal silhouette score occurs when the number of clusters is 3. (**g**) PCA projections of the phasic activity patterns for all units, colored by lower bound of respiratory coherence. Highlighted points in (g) correspond to the example units of the corresponding color in (a). (**h**) Clustering is repeated with multiple clustering and data processing techniques (see methods), for multiple coherence threshold values. The silhouette scores for those techniques are shown as a function of number of clusters. When weakly coherent units are included (coherence threshold =[0.1,0.3,0.5]), the optimal number of clusters is three (inspiratory, expiratory, tonic). However, if only strongly coherent units are included (coherence threshold = [0.7,0.9,0.99]) the optimal number of clusters is two (inspiratory, expiratory), as the tonic neurons have been excluded prior to clustering.

An example recording of 128 simultaneously recorded single units is shown in Fig. 1c, along with integrated diaphragm for 4 breaths. As expected, single units exhibit varied breath-averaged activity patterns (Fig. 1d). Since respiratory rate and inspiratory durations vary over time and across animals, we map each breath to phase (*ϕ*) such that the onset of inspiration occurs at *ϕ* = 0, inspiratory bursts have 0 ≤ *ϕ* ≤ *π*, and inter-breath intervals (referred to as expiration throughout) have −*π* < *ϕ* < 0. We avoid more granular discretization of phase (e.g., pre-inspiratory, which has no quantitatively defined temporal marker) in favor of reporting the exact *ϕ* values. We then quantify each neuron’s activity as a function of this normalized phase (Fig. 1e-h). Individual neurons are parameterized by coherence (quantifies how strongly a neuron shares frequency components with breathing, Extended Data Fig. 3), and phase lag (i.e., phase of breathing cycle in which the neurons activity is biased). We compute the coherence and phase lag with Chronux^44^(See Methods), and use the lower bound on coherence (*C_lb_* ; 99.9% confidence interval) so as to not categorize some neurons as more coherent than statistically likely. Neurons with *C_lb_* > 0.1 were categorized to be phasic, and any neuron with *C_lb_* ≤ 0.1 to be tonic; this categorization is used throughout. We choose a low value of *C_lb_* to include neurons that may be only weakly correlated with breathing (Extended Data Fig. 3b). Putative axonal units had lower respiratory coherence and firing rates than putative somatic units (Extended Data Fig. 2e). Slightly more than half of all somatic units showed evidence of respiratory coherence (3,637 of 6,315, 57%). Phasic neurons were more likely to be inspiratory (3,637 of 6,315; 60%) (Fig. 1gh). On average, inspiratory units have higher firing rates, but are not more coherent than expiratory units (Extended Data Fig. 3c,d,g).

**Figure 3.**
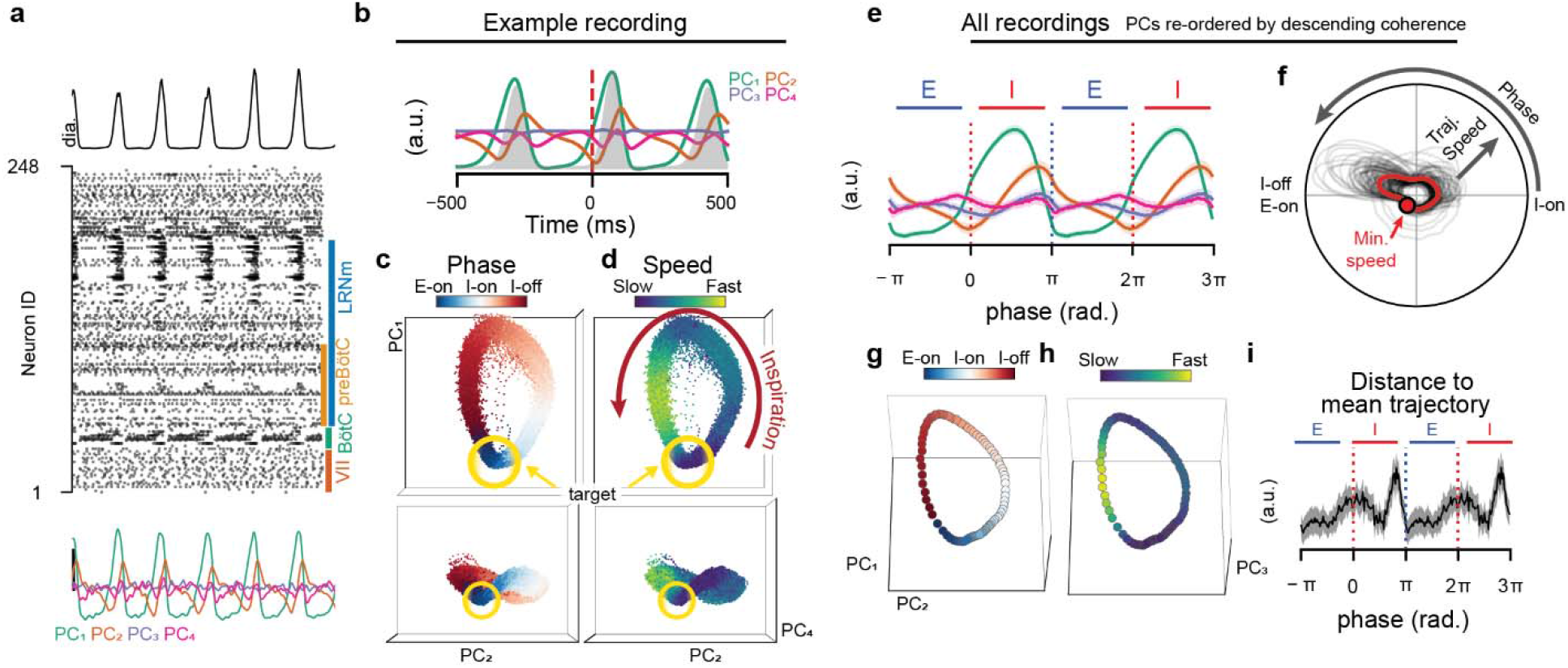
Low dimensional manifold underlie respiratory neural activity. (**a**) Example integrated diaphragm (top), raster (middle) for 248 simultaneously recorded units, and four leading principal components (bottom; green, orange, purple, pink, respectively) that decompose the neural activity. (**b**) Breath-aligned average PCs from (a). Mean diaphragm activity is shaded gray region. PC colors as in (a). (**c**) Projection of the neural population into the space of the PC1,PC2 and PC4 for 10 minutes of recording time. PC3 is not shown as it is not coherent with the breathing cycle for this recording. Each dot is the 3-dimensional position of these three PCs in a 5ms time bin. Color represents the breathing phase. Inspiration onset occurs where dots are white; offset occurs where dots transition from red to blue. (**d**) as in (c), but color represents the instantaneous temporal derivative of the trajectory through the PC space (**e**) After reordering the PCs by respiratory coherence (leading PC is most strongly coherent with respiration), we average the phase aligned values of the four most coherent PCs across all recordings. Zero and 2*π* are diaphragm onset, *π* and 3*π* are diaphragm offset. PC color as in (A). Shaded region is mean + S.E.M. (**f**) Angular plot of trajectory speed (radial axis) as a function of breathing phase (angular axis) for all recordings. Individual recordings in black, mean across recordings in red. Top half of circle is inspiration, bottom half is inter-breath interval. Minimum speed across all recordings is shown with a marker. (**g**) Average PC trajectory in the coherence re-ordered space across all recordings colored by phase as in (c). (**h**) same as (g) colored by trajectory speed. Each dot is a 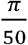 radian bin. (**i**) For each recording, mean trajectory through the PC space was computed. Then, for each breath, at each phase bin, the Euclidean distance to that mean trajectory was computed as a measure of breath-to-breath variability in latent trajectory. Distance to mean trajectory was then averaged across each breath and recording. High distance to the mean trajectory indicates high breath-to-breath variability at that phase of the breath.

**Figure 4.**
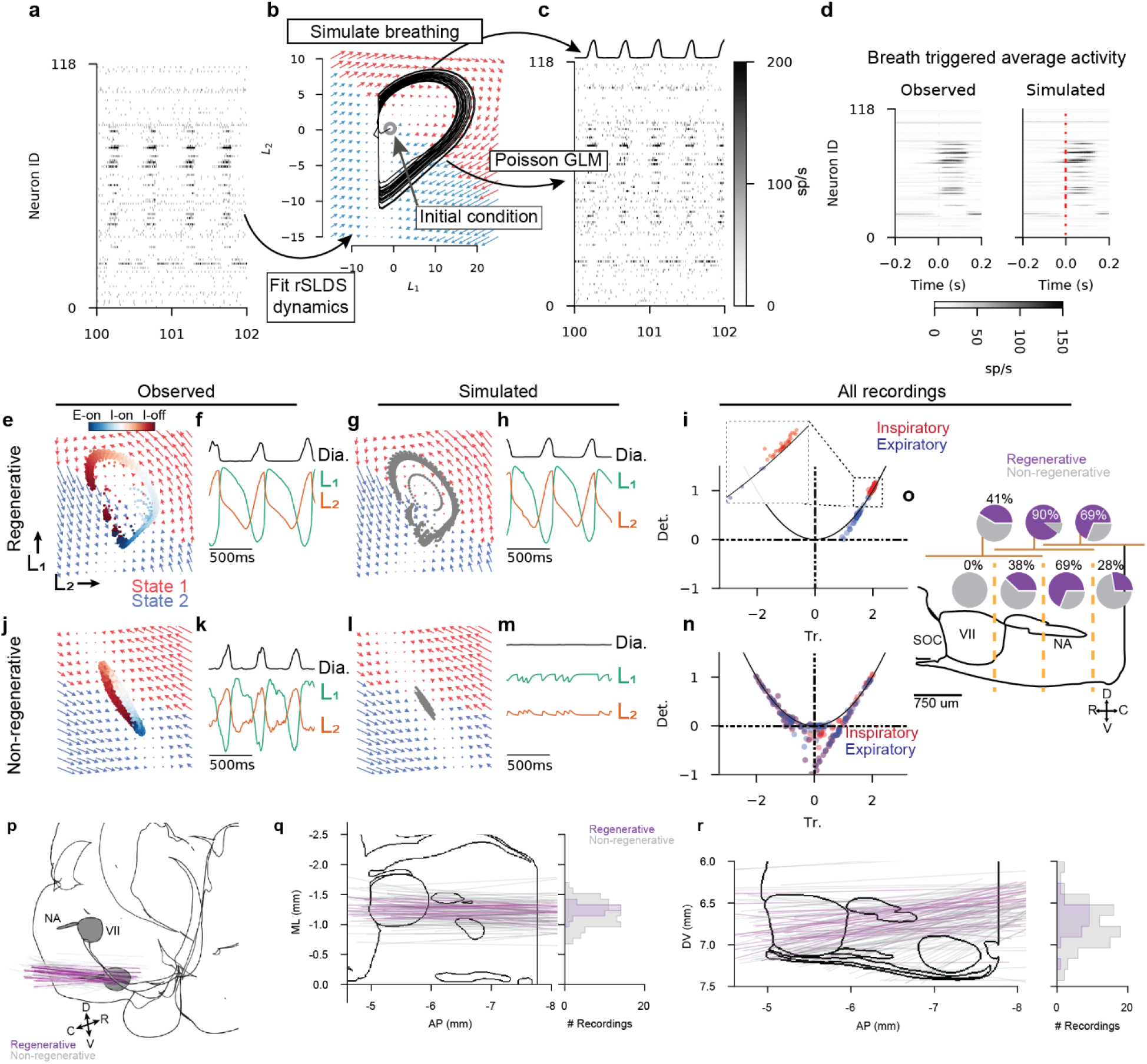
Inspiration and expiration follow alternating linear dynamical systems. (**a**) An example of two seconds of recorded neural activity across 118 VRC units. Color scale is shown in (c). (**b**) The rSLDS model fits a two-state dynamical system in two dimensions; the two flow-fields indicate these dynamics. Red arrows indicate state 1 dynamics, blue arrows indicate state 2 dynamics. The state that governs the evolution of the latent trajectory is determined by the current value of the latent. Trajectories can be simulated by supplying an initial state condition (0,0) and allowing the network to evolve according to the dynamics. (**c**) A Poisson GLM estimates the spike rate of each neuron based on the position of the system in the latent space as it evolves over time. A support vector regression simulates the diaphragm activity (top) based on the given latent state over time. (**d**) Peri-event time histograms (PETHs) of breath onset aligned spike rate averages for all 118 recorded neurons (left) and for simulated breath aligned spike rate averages for simulated neurons (right). Simulated PETHs match observed PETHs. (**e**) The observed trajectory through the 2D latent space is shown as dots (5ms time bins). (**f**) Observed diaphragm activity (black) and the two latent variables (green, orange) over ~1s. (**g**) The inferred dynamics (flow fields identical to (e)) are used to simulate evolution through the latent space. Gray dots indicate simulated latent. (**h**) as in (f) for simulated latent evolution. (**i**) Inferred latent dynamics were clearly active during inspiration or expiration. The trace (Tr.) and determinant (Det.) are shown for the dynamics matrices that govern either inspiration (red) or expiration (blue). Parabola is the equation 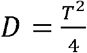. (**j-n**) as (**e-i**) for recordings that did not simulate rhythmic diaphragm activity. (**o**) regenerative rSLDS models were given access only to rostro-caudally restricted subsets of recorded units. Vertical yellow dashed lines indicate borders of the four anatomical subsets imposed. Pie charts indicate percentage of recordings that maintain regenerative capability (purple) with indicated subsets. Bottom row includes only one subset, top row includes units of two neighboring subsets indicated by dark yellow horizontal lines. (**p**) Three-dimensional and two-dimensional projections (**q,w**) of the probe tracks that resulted in regenerative (purple) or non-regenerative (gray) dynamics. Marginal histograms show the Mediolateral (q, right) and dorsoventral (r, right) location of the probe midpoint.

The phase-averaged and max rate normalized activities of all putatively somatic units with significant respiratory coherence smoothly tile the respiratory cycle (Fig. 1f). Notably, there is sequential recruitment of neurons during progression of the inspiratory (I) phase. While the inspiratory and expiratory phases recruit largely non-overlapping neural populations, the boundary from expiration to inspiration is straddled by many neurons. By contrast, fewer neurons straddle the inspiration to expiration transition (Fig. 1f). Thus, the onset of inspiration is a gradual, continuous neural process while the offset of inspiration is a sharp transition.

We next determined the anatomical location of each single unit. We reconstruct the 3D location of the fluorescent probe tract with respect to the AllenCCF and identify the location of the channel on which the neuron’s spike waveform was largest. Inspiratory and expiratory neurons were found continuously distributed throughout the VRC (Fig. 1i, Extended Data Fig. 3-6). As expected, proportionally more phasic neurons were found along the anatomical bounds of the VRC, while tonic neurons were more widely spatially distributed (Fig 1i, Extended Data Fig. 3-6, Extended Data Video 1). These distributions were not qualitatively affected by varying the coherence threshold (Extended Data Fig. 3a). However, as the coherence threshold is raised (i.e., we consider only strongly coherent neurons in the analysis), classical parcellations of the VRC become more pronounced (Extended Data Fig. 3, Extended Data Fig. 5). In all locations, inspiratory neurons were more numerous than expiratory neurons, and there is a predominant rostro-caudal gradient where expiratory neurons are found rostrally and inspiratory neurons are found caudally (Extended Data Fig. 3-5).

**Figure 5.**
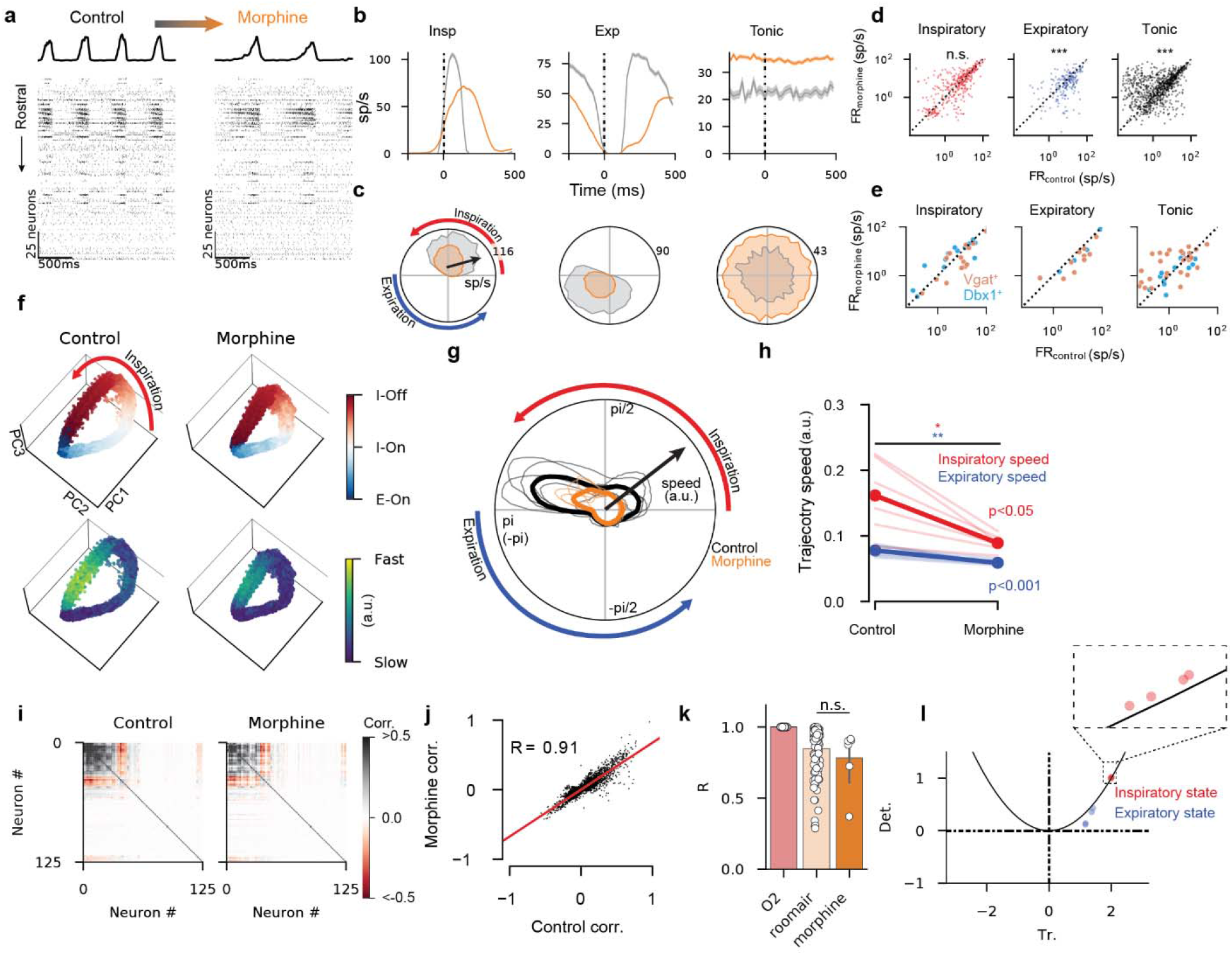
Opioids cause temporal restructuring of respiratory neural activity, but do not alter underlying manifold. For all panels, gray is control, orange is morphine. (**a**) Example rasters and diaphragm activity before (left) and after (right) IP administration of 150mg/kg morphine. (**b**) Breath onset and (**c**) phase aligned average activity of three example units in control and after morphine administration. Top half of circle is inspiration, bottom half is expiration. Mean +/− S.E.M. is shown. (**d**) Firing rates of inspiratory (red, n=386), expiratory (blue, n=217) and tonic (black, n=894) units before (x-axis) and after (y-axis) morphine administration. (*** p<0.0001, one sample t-test of log transformed ratio of firing rates) (**e**) Firing rates of optogenetically tagged units (blue: Dbx1^+^, orange: Vgat^+^) separated by respiratory firing pattern. (**f**) Low-dimensional population projections onto the leading three principal components. Each dot is a 5ms bin. Top row color indicates phase, bottom row color indicates trajectory speed. (**g**) Angular plot of trajectory speed (radial axis) as a function of breathing phase (angular axis) for all recordings. Control is black, morphine is orange. Individual recordings are thin traces, mean across all recordings are thick traces. (**h**) Average speed during inspiratory phase (red) or expiratory phase (blue) in control vs morphine (2 sided paired t-test *p<0.05, **p<0.001). Individual recordings are thin traces, means are thick traces. Trajectory speed units are arbitrary. (**i**) Correlation of each pair of recorded units in control (left) and morphine (right), ordered by ward clustering. (**j**) Correlation value of each pair of units before (x-axis) and after (y-axis) morphine administration. Each dot is a pair of units. (**k**) Linear fit (R-value) of the pairwise correlation values in O_2_ compared to room air and during morphine administration. We consider room air as a control change in correlation structure as compared to O_2_. Changes in population correlation structure after morphine administration are not different than those observed in room air (Mann-Whitney U-test p=0.1). (**l**) Trace-determinant plane for the dynamics matrices that govern the inspiratory and expiratory states in the rSLDS models fit during OIRD. Only recordings in which the dynamics were regenerative during control eupnea (n=4) are shown. Parabola is the equation 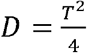.

Lastly, we assessed the respiratory related neural activity patterns and anatomical distribution of optogenetically identified, somatic units. Units of all optogenetic identities and phasic firing patterns are found widely distributed throughout the VRC (Fig. 1j-l). However, some populations over represent certain phases of breathing (Fig. 1l, Extended Data Fig. 6ef) On average, Vgat^+^ and Dbx1^+^ neurons are more strongly respiratory tuned than the Vglut2^+^, ChAT^+^, or untagged populations (Fig. 1m). Vgat^+^ neurons over-represent expiratory phase; and inhibitory, expiratory neurons tend to be—but are not exclusively—found in the rostral VRC (i.e., BötC/pFRG). Dbx1^+^ neurons overrepresent the inspiratory phase and these inspiratory neurons are found more caudally in the VRC.

### Activity based cell clustering reveals continuously distributed patterns

VRC units are typically categorized into one of several groups based on their phase averaged activity pattern. These categories describe the relationship between the unit’s activity and phase, and allow for comparisons across units. For example, a typical pre-inspiratory unit will exhibit a ramp in firing-rate prior to the onset of the breath (or burst *in-vitro*) and decay during the inspiratory effort (Fig. 2a). While many studies agree qualitatively on inclusion of units into these categories^7,45,46^, the number of categories varies across studies, and no quantitative categorization scheme exists. Several units that exemplify some of these categories are shown in Fig 2a.

The ability to record from thousands of respiratory neurons affords data-driven clustering methods for identifying groups of neurons that exhibit similar phase aligned activity. We first normalize the binned (100 bins, phase-averaged activity from −*π* to *π*) of each somatic unit between zero and one, and then compute the principal components across all units to reduce the dimensionality for subsequent clustering. We perform both hierarchical clustering and k-means clustering on both the full data (each unit has 100 features, one for each phase bin) and the reduced data (each unit has 4 features, one for each retained PC, >95% variance explained). We vary the number of discovered clusters from 2 to 15 to determine the optimal number of activity-based clusters as defined by maximal silhouette score.

We find that splitting into three clusters returns inspiratory, expiratory, and weakly-phasic cluster means; splitting into six clusters returns cluster means consistent with classical categories; and splitting into 15 clusters returns cluster means that sequentially tile phase (Fig 2b-e). We compute the silhouette score and find that the optimal number of clusters is three (Fig. 2f,h), and that further discretization over-splits the data. However, we include even weakly coherent units in these analyses which could contaminate the clustering and give an erroneous appearance of continuity in the population (Fig. 2g). Therefore, we systematically vary a coherence threshold such that we cluster only on increasingly exclusive populations of increasingly coherent units. We find that as weakly coherent units are excluded, the maximal silhouette score becomes two, and the cluster means resemble inspiratory and expiratory units only (Fig. 2h).

### Respiratory neural populations are restricted to trajectories on a low-dimensional manifold that target the offset of inspiration

Recording of simultaneous neural activity allows for the quantification of low-dimensional, coordinated activity patterns across the population of recorded neurons. The low-dimensional subspace that contains the majority of the neural variance is referred to as a neural manifold^3,47^, while the evolution of the low-dimensional activity through that subspace is referred to as its latent dynamics. We first use principal components analysis (PCA) to decompose the high-dimensional activity of simultaneously recorded units (Fig. 2a-d, Extended Data Fig. 7, both putative axonal and somatic units are included). Low dimensional neural activity evolves through a consistent rotational trajectory in the principal component (PC) space, and the location along that trajectory is highly correlated with the respiratory phase (Fig. 2b,c,e,g, Extended Data Videos 2,3). These trajectories show conserved temporal evolution such that the trajectory is fastest (i.e., the population activity changes most quickly) during the inspiration-off/expiration-on transition (Fig. 2d,h). To compare PCs across recordings, we re-order the PCs by coherence such that the leading PC is the most coherent with respiration, regardless of the variance accounted for (Extended Data Fig. 7). We compute the average low-dimensional trajectory across all recordings (Fig. 2e-i) by first re-ordering the PCs by their respiratory coherence, and then normalizing the time-domain to phase. Ordered PC_1_ exhibits pre-inspiratory and inspiratory activity, PC_2_ augments during inspiration and peaks at the offset of inspiration, and PC_3,4_ exhibit more complex, but consistent, relationships with phase. Both trajectory speed (which quantifies rate of change of neural state) and distance to mean trajectory (which quantifies the breath-to-breath variability of neural state) are maximal prior to offset of inspiration, and minimal just after offset of inspiration (Fig. 2f,i). This suggests that the peak of inspiration represents an unstable region of the latent space, and the offset of inspiration resembles an attractive “target”. This low-dimensional landscape is conserved across recordings and animals (Extended Data Fig. 7).

### Alternation between discrete, unstable dynamical landscapes for inspiration and expiration recapitulate VRC population activities

The above PCA analysis inherently treats all points in time as independent and does not incorporate temporal dynamics of the neural activity. Therefore, we estimate latent linear dynamics from the high-dimensional observations (i.e., the activity of the recorded neural population) using a recently developed recurrent switching linear dynamical system (rSLDS)^48^. This model allows for multiple sets of linear dynamical systems to govern the latent evolution (switching). The dynamics to be employed at any one point in time are determined by the current latent state (recurrent).

We fit 2-state, 2-dimensional rSLDS (see Methods) to compute the latent dynamics of the neural population (Fig. 3a,b, Extended Data Fig. 8, both putative axonal and somatic units are included). We then simulated the latent dynamics (Fig. 3b) and resultant neural activity (Fig. 3c) by supplying the dynamics matrices as inferred from the experimental data and visualized by the flow fields in Fig. 3b. Simulated firing rates were estimated using a Poisson GLM with the latent state as inputs. Lastly, we use support vector regression to predict the diaphragm activity from the two-dimensional latent state (Fig. 3c). This allows us to simulate the entire VRC population and diaphragm activity from only the two-dimensional latent state. By aligning the simulated firing rates to the simulated diaphragm onsets, we can well recreate the breath-averaged activity of the recorded VRC population (Fig. 3d). We found that the dynamics fit from a recording can either be “regenerative” (Fig. 3e-i), in that the dynamic matrices (i.e., flow-fields) alone are sufficient to recreate oscillatory latent state evolution and subsequent rhythmic simulated diaphragm activity (Fig. 3gh), or non-regenerative (Fig. 3j-m). In the non-regenerative case, simulations could not produce oscillations or rhythmic simulated diaphragm activity, despite oscillations in the observed latent dynamics (Fig. 3jk). If a model was non-regenerative, it means that the neural population sampled was not sufficient to generate oscillatory latent patterns. Models with only 1 state (K=1, i.e., not switching^48^) could never be regenerative (Extended Data Fig. 8), but rather formed decaying spirals; models with three states were scarcely more likely to be regenerative than models with two states (n=35/116 regenerative for K=2, 37/116 regenerative for K=3; Extended Data Fig. 8).

We then investigate the properties of the dynamics matrices as inferred by the rSLDS models. The two inferred states are clearly restricted to either the inspiratory or expiratory phase of the breath. That is, the neural population follows one set of dynamics during inspiration, and another during expiration. For the regenerative recordings, the dynamics were strikingly similar across recordings (Extended Data Fig. 8d). By examining the trace-determinant plane for the Jacobian of each state (Fig. 3i), we find that inspiratory dynamics were governed by spiral sources 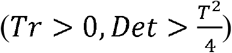, while expiratory dynamics were governed by nodal sources 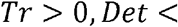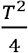. These unstable regimes intersect to generate alternation between these two states.

To test whether rostro-caudal subregions of the VRC were sufficient to recapitulate oscillatory dynamics, we re-fit and re-simulated all the regenerative recordings but systematically removed subsets of the population along the rostro-caudal extent of the recording. Models with access to the caudal or rostral extremes of the VRC were less likely to be regenerative than those with access to the middle sections which contain the preBötC and BötC (Fig. 3o).

In contrast, non-regenerative recordings showed no clear patterns in their dynamics across recordings and states (Fig. 3n, Extended Data Fig. 8e). Likely, recordings that were non-regenerative did not sample the appropriate neural populations to adequately reconstruct rotational dynamics. We found that whether the recording was regenerative or not depended primarily on the mediolateral anatomical location of the recording. If the recording was restricted to the center of the VRC, where many respiratory tuned neurons are located, it was more likely to be regenerative than at the borders of the VRC (Fig. 3p-r).

### Opioids alter timing, but not latent structure, of neural population activity

We next sought to quantify network-wide activity changes in response to systemic opioid administration. To investigate network-wide neural reconfiguration during opioid-induced respiratory depression (OIRD), we administered a single dose of morphine (150mg/kg, 4 Vgat^Cre^;Ai32 mice, 2 Dbx1^CreERT2^;Ai32 mice) and recorded from the VRC. Respiratory rate decreased by an average of 21% (± 9% S.D), and inspiratory duration increased by an average of 105% (± 22% S.D) (Extended Data Fig. 9). An example raster of 175 simultaneously recorded neurons and associated diaphragm activity is shown before and after morphine administration (Fig. 5a). Gas supplied was 100% O_2_, and we include both somatic and axonal units in these analyses.

We first determined the extent to which morphine altered the activity of single neurons of the VRC. We found that morphine does not unilaterally reduce the firing rate of neurons, but instead induced coordinated changes across the entire population of VRC neurons (Fig. 5b-e). The duration of spiking in individual neurons increased and spike rates changed for many neurons, but the phasic patterns were preserved (Fig. 5bc, Extended Data Fig. 9). Inspiratory, expiratory, and tonic neurons all show variable changes in firing rate after morphine administration (Fig. 5de). However, on average, expiratory neurons have reduced firing rate in morphine, inspiratory neurons are unchanged, and tonic neurons have increased firing rates (Fig. 5de, Extended Data Fig. 9d). Of the 1,497 units recorded after morphine administration, 56 were identified as Vgat^+^ (inhibitory)and 31 as Dbx1^+^ (excitatory). Both Vgat^+^ and Dbx1^+^ expiratory units have reduced firing rates, while the Vgat^+^ tonic units showed increased firing rates (Extended Data Fig. 9j,k).

To determine the extent to which coordinated population activity changes during OIRD, we examined the neural manifolds before and after morphine administration. We found that PC trajectories of the neural population were qualitatively similar across conditions (Fig. 5f, Extended Data Fig. 9g, Extended Data Fig. 10). The relationship between location in the PC space and breathing phase was conserved (Fig. 5f), indicating that the underlying latent structure of the population remained largely unchanged. The inspiration-off target was preserved during opioid administration (Fig. 5f). Surprisingly, morphine slowed the evolution of the trajectory through PC space (Fig. 5g, Extended Data Video 4). This slowing occured at most phases of the respiratory cycle, but most dramatically during the second half of the inspiratory burst (Fig. 5gh). By examining the breath aligned averages of the PCs we saw that morphine slowed the latent temporal evolution of the PCs, but the phasic relationship between these PCs was preserved.

To confirm that relations between neural activity patterns are preserved during OIRD, we computed the pairwise zero-lag correlations of spike rates for all neurons in baseline (100% O_2_) and after morphine administration. We found that the change in correlation structure during OIRD was not significantly different than the change observed when exposed to room air (Fig. 5i-k). We further quantified the changes in low-dimensional structure by separately computing the PCA decompositions in control and in morphine, and then comparing the subspaces spanned by the leading eigenvectors of those decompositions^49^. We quantified the similarity between these subspaces using principal angles^50^ (see Methods) and found that the subspaces largely overlapped, indicating conservation of the low-dimensional population structure in OIRD (Extended Data Fig. 13). Lastly, to confirm that the latent dynamical structure was not disrupted during OIRD, we re-fit the rSLDS model only on data recorded after morphine administration, for the recordings that were regenerative during normal breathing (n=4). In recordings that were regenerative during control conditions we found that spiral sources and spiral nodes continued to govern inspiration and expiration, respectively. (Fig. 5l, Extended Data Fig. 14).

### Sighs are low-dimensional excursions during inspiration which disrupt the following breath

Sighs are intermittent, large amplitude breaths that occur at regular intervals^51–53^. Sighs are critical for maintaining blood gas homeostasis by preventing atelectasis; loss of this behavior is fatal^54^. As sighs are known to cause reconfiguration of VRC activity and are triggered by activity from the RTN/pFRG^55^, we sought to determine how VRC population activity was modulated during sighing.

A sigh is exemplified by increased diaphragmatic activation, and a prolonged period of apnea after the sigh (Extended Data Fig. 11). In this experimental preparation we could evoke sighs by presenting room air (21% O_2_) instead of 100% O_2_ (Fig. 6a). Sighs show pronounced increase in firing rates of inspiratory neurons throughout the VRC (Fig. 6b-d, Extended Data Fig. 11).

**Figure 6.**
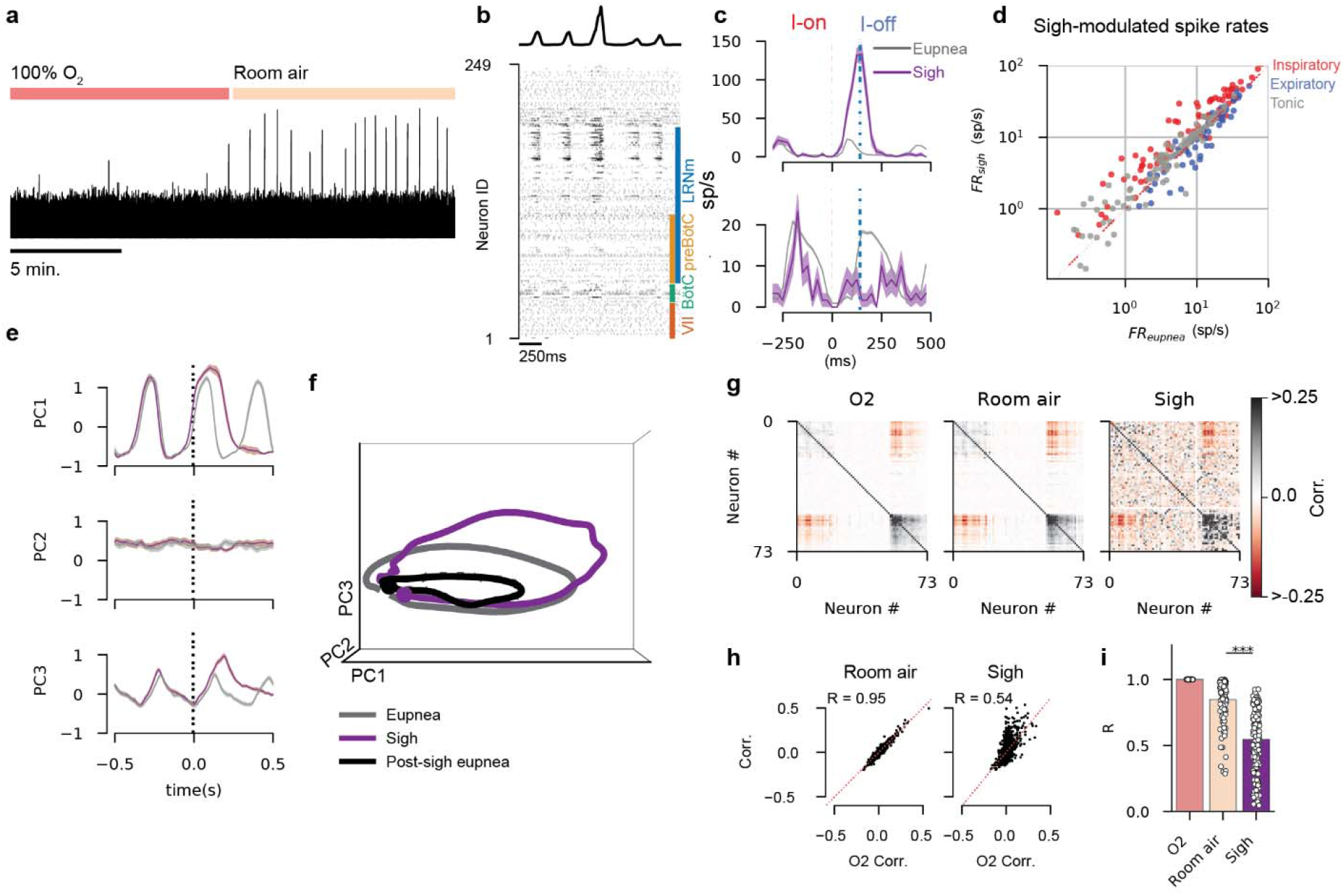
Sighs are inspiratory excursions that disrupt subsequent breaths. (**a**) Integrated diaphragm trace over 20 minutes show increased frequency of sighs during room air presentation. (**b**) Example spike raster and diaphragm activity before and after a sigh. (**c**) Breath onset aligned average activity of two example units (mean F.R. +/− S.E.M.) for eupnea (gray) and sighs (purple). Firing rate of top unit increases, and bottom unit decreases during the sigh. (**d**) Firing rates of all units recorded in (b) for eupnea (x-axis) and sighs (y-axis). Each dot is a recorded unit. Color indicates preferred phase of firing during eupnea: inspiratory (red), expiratory (blue) or tonic (gray). (**e**) Breath onset aligned average of the leading three principal components in eupnea (gray) and sighs (purple) for an example recording (**f**) Average population trajectories through PC space for the recording in (e) for eupnea (gray), sighs (purple), and the breath after a sigh (black). (**g**) Correlation values for all pairs of units during eupnea in O_2_, room air, and during sighs. Red is strong negative correlations; black is strong positive correlations. (**h**) Correlation value of each pair of units during O_2_ presentation (x-axis) against those during room air presentation (y-axis, left) and during sighs (y-axis, right). (**i**) Linear fit (R-value) of the pairwise correlation values in O_2_ compared to room air and sighs. We consider room air as a control change in correlation structure as compared to O_2_. Changes in population correlation structure during sigh are different than those observed in room air (***Mann-Whitney U-test p<0.001).

Examination of the low-dimensional trajectories of eupnea and sighs shows that sigh trajectories overlap with eupnea, exhibit an augmented excursion during the inspiratory phase, and target the same post-inspiratory region that was observed during eupnea (Fig. 6ef, Extended Data Video 2) This altered trajectory is consistent across animals and recordings (Extended Data Fig. 11f). The trajectories of the subsequent breath after the sigh trace out the same shape as a eupnea in the PC space, but with smaller radii, indicating the neural population is incompletely recruited in the breaths following a sigh (Fig. 6f). The recovery to normal eupnea after a sigh takes several breaths (Extended Data Fig. 11e).

We show that correlated activity across the VRC population is altered during sighs by comparing the pairwise correlations between eupnea in 100% O_2_ with eupnea in room air and with sighs (Fig. 6g-i). The pairwise correlations are altered during sighs in comparison to eupnea in room air, but the principal angles between the eupneic and sigh indicate that the subspaces are largely overlapping (Extended Data Fig. 13), indicating a minor and temporary disruption of VRC manifolds.

### Gasps collapse rotational dynamics into ballistic trajectories

Under severely hypoxic conditions, the respiratory network generates auto-resuscitative gasps^36,37,56^. During gasping, breaths become punctuated large amplitude events with long inter-gasp intervals, larger diaphragmatic activity, and recruitment of more auxiliary inspiratory muscles (Fig. 7a)^37^. Inspiratory, expiratory, and tonic neurons of all genotypes have reduced firing rates on average (Fig. 7b-d, Extended Data Fig. 12a-c). Activity during the inter-burst interval is reduced across the population as evidenced by an increase in respiratory tuning strength (Extended Data Fig. 12f). Many normally expiratory neurons reconfigure and become active during inspiration (Extended Data Fig. 12de).

**Figure 7.**
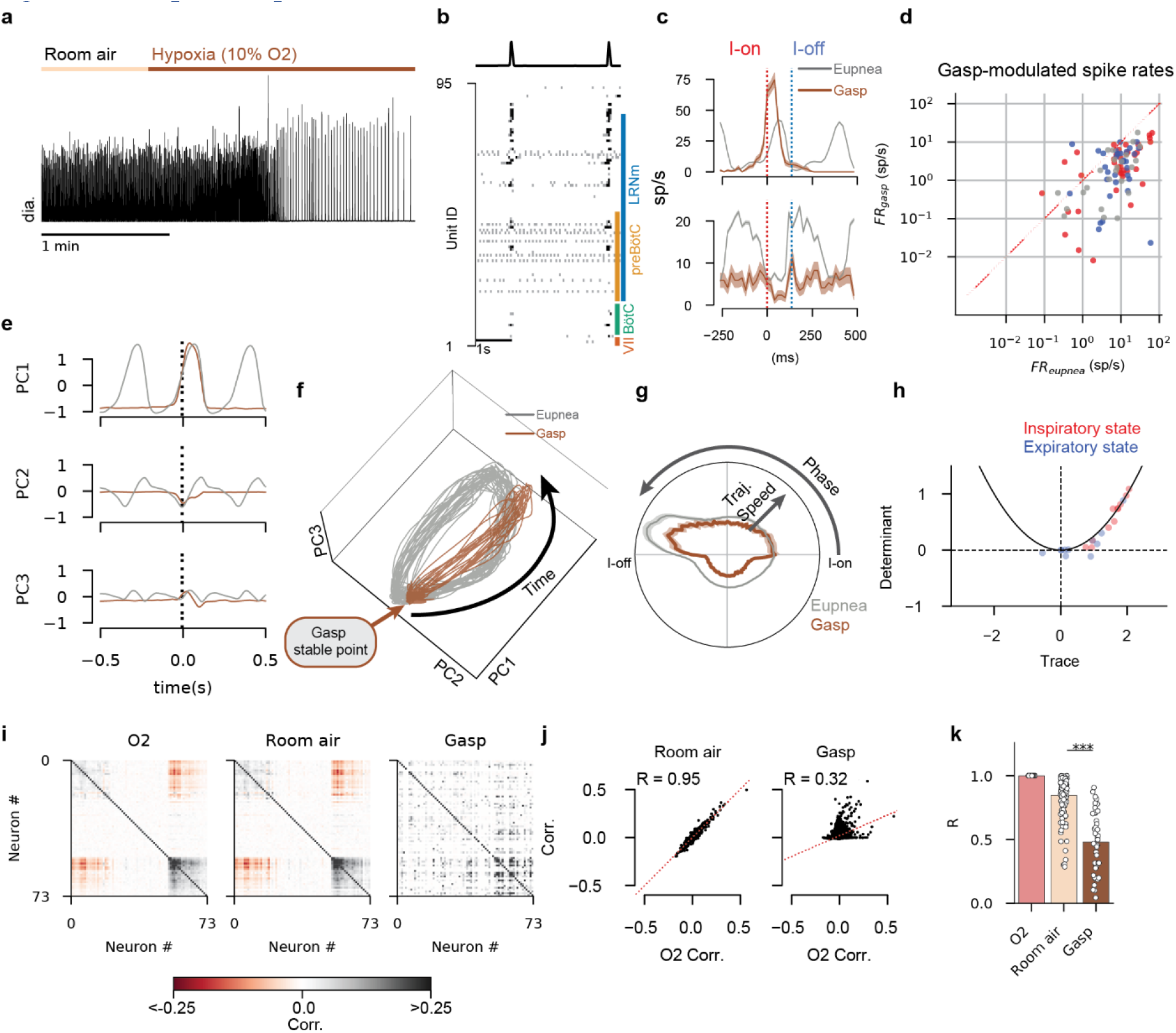
Gasps collapse latent orbits into ballistic, all-or-none efforts. (**a**) Integrated diaphragm activity illustrating the transition to gasping in during hypoxia presentation. (**b**) Example spike raster and diaphragm activity during a period of gasping. (**c**) Breath onset aligned average firing rates of two example units. Firing rate of top unit increases, and bottom unit decreases during gasping. Shaded region is mean +/− S.E.M. (**d**) Firing rates of all units recorded in (**b**) for eupnea (x-axis) and gasps (y-axis). Each dot is a recorded unit. Color indicates preferred phase of firing during eupnea: inspiratory (red), expiratory (blue) or tonic (gray). (**e**) Breath onset aligned average of the leading three principal components in eupnea (gray) and gasps (brown) for the recording in (b). (**f**) Population trajectories through PC space for the recording in (b) for eupnea (gray) and gasps (brown). (**g**) Trajectory speed (radial axis, a.u.) as a function of breathing phase (angular axis) for eupnea (gray), and gasps (brown). Top of circle is inspiratory effort, bottom of circle is inter-breath interval. (**h**) Trace-determinant plane for the dynamics matrices that govern the inspiratory and expiratory states in the rSLDS models fit during gasping. Only recordings in which the dynamics were regenerative during control eupnea (n=10) are shown. Parabola is the equation 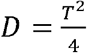. (**i**) Correlation values for all pairs of units during eupnea in O_2_, room air, and during gasps. Red is strong negative correlations; black is strong positive correlations. (**j**) Correlation value of each pair of units during O_2_ presentation (x-axis) against those during room air presentation (y-axis, left) and during gasps (y-axis, right) (**k**) Linear fit (R-value) of the pairwise correlation values in O_2_ compared to room air and gasps. We consider room air as a control change in correlation structure as compared to O_2_. Changes in population correlation structure during sigh are different than those observed in room air (***Mann-Whitney U-test p<0.001).

Importantly, the rotational dynamics and inspiratory-off target observed during eupnea are lost during gasping. Instead, the low-dimensional trajectories reconfigure to exhibit all-or-none, ballistic dynamics in which both pre-inspiratory and post-inspiratory activity is lost, and the PCs evolve only during the gasp effort. (Fig 7e-g). During gasping, the population is constrained to a stable region between gasp efforts, and each gasp is characterized by a quick and symmetric excursion away from, and back to, this region (Fig. 7e-g, Extended Data Video 5). Further, the transition from eupnea into gasping is smooth, characterized by a progressive “shrinking” of the rotational trajectories (Extended Data Fig. 12i). To examine the structure of the latent dynamics during gasping, we re-fit the rSLDS model to the period of time in which the animal was gasping, and only for the recordings that were regenerative during normal breathing (n=10). We find that the fixed points no longer cleanly separate into separate dynamics for inspiration and expiration as they did during eupnea (Fig. 7h, Extended Data Fig. 14). Lastly, we quantify the changes in correlation structure during gasping as was done for morphine and sighs (Fig. 7i-k). The correlation structure is drastically altered during gasping across all recordings, indicative of a VRC wide reconfiguration.

## Discussion

Growing adoption of large-scale population recording and analysis techniques has driven widespread appreciation of low-dimensional patterns as a ubiquitous feature of neural circuits^49,57^. Here, we show that this framework elucidates the network dynamics of distributed VRC populations that are critical to generating, patterning, and maintaining both normal and disrupted breathing. The VRC population evolves through elliptical trajectories on a low-dimensional manifold of neural activity space that are consistent across individual animals. These trajectories target an “inspiratory-off” attractive region, expanding the well-described phenomenon of an inspiratory off-switch^27–29^ to the behavior of the entire VRC. Further, VRC populations can be fully simulated by a simple dynamical model in which inspiration and expiration respectively are governed by alternation between discrete, intersecting, unstable dynamical regimes.

The high-resolution mapping data support the general organizational structure of the VRC as proposed in previous models^6,7^, but also reveal that these models are under-resolved to fully capture the distributed, heterogenous nature of the neural populations that underlie breathing and its associated behaviors. Indeed, our data further supports the idea that discrete boundaries between VRC subdivisions are not apparent in freely breathing animals^9,58^. We emphasize that these conclusions do not contradict the fact that the preBötC is necessary for breathing when lesioned acutely^36,59,60^ or when isolated in slices^13^, nor that the BötC is a region within the VRC that is biased towards inhibition of inspiration^6^.

This work is a significant advance on previous efforts to create maps of VRC activity patterns. Recently, Dhingra et al.^61^ performed a comprehensive mapping of local field potentials across the isolated rat brainstem; and the large body of work from Morris, Lindsay, and colleagues^46,62^ also recorded from many isolated respiratory units simultaneously in cats. The data presented here is restricted to the VRC but offers advances on those works in that we record simultaneously from an order of magnitude more single units in animals with a fully intact neuro-respiratory system breathing in a dynamic regime similar to that during awake, quiescent breathing. Further, we employ optogenetic tagging to identify neurons that express particular transcriptional markers (e.g., excitatory vs. inhibitory). Importantly, we find that the proliferation of functional cell classes they and others describe, at least at the level of the VRC, are better quantitatively described as a continuum of activity patterns, rather than sets of discrete functional types.

A major advantage afforded by large scale recording and subsequent low-dimensional analyses is that it allows intuitive interpretation of network-wide compensations to perturbations that would otherwise be impenetrable. Systemic opioid administration evokes widespread and diverse changes in single cell spiking activity. However, the low-dimensional rotational behavior of the VRC is largely preserved during OIRD, suggesting the possibility of compensatory network solutions to maintain robust breathing. The overarching effect of OIRD can be described as a slowing or “frictional” effect on the network dynamics. Opioids have been demonstrated to cause both hyperpolarization of Oprm1^+^ neurons, as well decreases in synaptic efficiency^32^. These mechanisms may, at the level of the network dynamics, impose a “dampening effect”, which in the extreme, may sufficiently slow breathing to result in inescapable hypoxia.

Subsequently, hypoxia induced gasping results in a collapse of the rotational dynamics. This collapse implies that network mechanisms that support normal breathing can no longer be maintained. Given that the neural population collapse into gasping is continuous, it may be the case that there is not a discrete “switch” or triggering mechanism involved in evoking gasping. Instead, gasping may come about as a degeneration of the normal eupneic behavior as it gradually becomes unable to be sustained by network interactions. These data augment observations in slices that show the cellular and molecular mechanisms governing rhythmic activity during eupnea are fundamentally different than that of gasping^56^; the latter is selectively dependent on serotonergic and noradrenergic mechanisms^63–66^. These modulatory systems have been found to be disrupted in children that died of SUIDS^38–40^. Importantly, the serotonergic disturbances are widely distributed in the medulla and include the nucleus gigantocellularis located in the rostral medulla, an area which is critical for arousal^67–70^. We speculate that these disturbances remain undetectable under normal breathing conditions because the VRC-wide rotational coordination between inspiratory and expiratory populations imbues the respiratory network with increased robustness. However, during hypoxic and gasping bouts, when the network dynamics transition from rotational to ballistic dynamics along the entire VRC, these rostrally located disturbances may become fatal in children susceptible to SUIDS.

The large-scale neural population dynamics associated with behaviors so stereotyped as breathing are just beginning to be explored^4,71^. The consistency of low-dimensional structures across animals seen here, and the striking similarity to the neural rotations in the spinal cord described by^4^ suggest that stereotypical behaviors may be governed by fundamental neural dynamics that are similar across motor behaviors and species. The similar low-dimensional structure of neuronal population dynamics contrasts with the striking cellular diversity and drastic differences in the biological substrate responsible for the generation and reconfiguration of these rhythmic behaviors. This suggests the possibility of a convergent and/or degenerate solutions for neural systems to perform computationally similar operations in disparate systems^5^.

## Data Availability

Data are available upon reasonable request

## Code availability

Custom MATLAB and python code used to process and visualize the data in this study are available upon reasonable request

## Acknowledgements

We would like to thank Lely Quina for contributions to histological processing, and Dr. Aguan Wei and Dr. Joshua Glaser for discussions and comments on the manuscript. We acknowledge Research Scientific Computing at Seattle Children’s Research Institute for providing high performance computing (HPC) resources that have contributed to the research results reported within this paper.

## Online Methods

### Animals

All procedures were approved by the Seattle Children’s Research Institute Institutional Animal Care and Use Committee. Both male and female homozygous Vglut2^Cre^ (Jax# 028863), Vgat^Cre^ (Jax# 016962), ChAT^Cre^ (Jax# 031661), and Dbx1^CreERT2^ (Donated by Dr. Del Negro College of William and Mary, VA) mice were crossed with homozygous Ai32 mice (Jax# 012569) which contain a flox-STOP-flox sequence fused to channelrhodopsin (ChR2) and enhanced yellow fluorescent protein (EYFP) at the Rosa26 locus. Thus, offspring expressed channelrhodopsin and EYFP only in the neurons that expressed the promoter for the Cre driver. In Dbx1^CreERT2^ animals, tamoxifen (24mg/kg IP) was injected at E10.5 to target neurons of the preBötC^11^.

### in-vivo surgical preparation and data acquisition

Anesthesia was induced in 3% isoflurane and urethane (150g/kg) was administered IP; animals were then removed from isoflurane. Animals were placed on a temperature regulating heating bed (Kent Scientific) and maintained at 37C° for the remainder of the experiment. Two fine (0.005 in. diameter) stainless steel wire EMGs (AM-systems) were inserted into the right diaphragm. The EMG signal was amplified (10,000x) and filtered (100 Hz to 5kHz) (AM-systems 1700) before being digitized at 10kHz (NI PXIe-8381). Ear bars were used to affix the head in a stereotactic frame. The skin overlying the scalp and neck musculature was removed and the skull surface cleared (Extended Data Fig. 1). The neck musculature was removed to expose the occipital bone, dural surface just ventral to the cerebellum, and the first cervical vertebra. The skull was then leveled such that bregma and lambda lie in the horizontal plane. A custom titanium headplate was then affixed to the skull surface with UV cure dental cement. The animal was removed from the stereotactic frame and re-positioned in a headfixing frame to allow for access to the nostrils while maintaining stereotactic level. A custom 3D printed nosecone which allowed for nasal pressure measurement as well as gas presentation was placed on the nose of the animal. 100% O_2_ was then supplied to the mouse. We then carefully removed the dura overlying the left caudal surface of the brainstem with a #11 scalpel, taking care not to damage underlying brain tissue or vasculature. A single Neuropixel probe (IMEC) was dipped in DiI (Thermofisher #V22885) and then placed in a motorized manipulator (Sutter MPC-325) positioned caudal to the animal. The probe holder was placed in the horizontal plane, with the probe tip toward the animal. We then find what we term the “skull apex” point (Extended Data Fig. 1), the point at the midline of the suture between the intraparietal and the occipital bone. We then target the VRC at 1.25mm lateral, 4.8mm ventral to this point. The probe is advanced to touch the caudal surface of the brainstem and the rostro-caudal position is recorded. The probe is then advanced 4mm into the brainstem at a rate of approximately 1mm/min. The probe settles for 15 minutes before recording begins. Neural and EMG data are acquired with SpikeGLX. Gas presentation was controlled by electrical solenoid valves (Beduan) connected to a single flowmeter (Cole Parmer). Input pressures were calibrated so that all gasses were presented at 150ml/min. Timing of solenoid valves and laser pulses (see Optical Tagging) were controlled by a Teensy 3.2 microcontroller. In some recordings, a single dose of morphine (150mg/kg) was administered IP following a baseline recording period (3-5min) and the optical tagging protocol. After recording, the probe is retracted, and repositioned approximately 100 m away in either the medio-lateral or dorso-ventral axis before another insertion and recording procedure. The probe is not redipped in DiI between each insertion. Following all recordings, the probe is extracted and placed in 1% Terg-a-zyme (Sigma-Aldrich #Z273287) overnight. The animal is perfused with saline and 4% paraformaldehyde, and the brain is removed for histological processing.

### Optical tagging

During recording, a Cobalt 473nm laser is attached to a 600*μ*m diameter 0.22NA fiber optic with a cleaved end (Doric). Fiber power measured at the tip was 50-70mW; high power and large fiber diameter was used to increase the likelihood of evoking spiking in rostrally located neurons that are far from the caudal surface of the brainstem. The fiber optic was placed just caudal to the brainstem surface. To perform optical tagging, 75 sequential 10ms laser pulses with a 1.5ms duration sigmoidal on-ramp and off-ramp and a 3 second delay between each pulse was presented to the caudal surface of the brainstem (Extended Data Fig. 1). This causes light-evoked spiking in neurons that express ChR2. This stimulation can directly alter respiratory activity and so periods of opto-tagging are not used in analyses. Positively tagged neurons are determined using the Stimulus Associated Latency Test (SALT^24^). Neurons with a SALT p<0.0001 and at least 25 optically evoked spikes were considered positively tagged.

### Data processing and spike sorting

The action potential band (acquired at 30 kHz) Neuropixels data were filtered (CatGT global demux), and spikes were extracted and sorted using Kilosort3 (see Table 1 for KS3 parameters). Putative single units were then quality sorted using the ecephys pipeline^72^. Double counted spikes were removed. All clusters were saved for potential reanalysis. Only units that were classified by the ecephys pipeline as not “noise” units, had ISI violations <2, amplitude cutoff <0.1, and a presence ratio>0.9, and were also labeled as “good” by KS3 were retained for analysis. This restrictive filtering potentially missed viable single units, but likely rejected most false positive units^72^.

Raw diaphragm EMG data (acquired at 10kHz) was processed offline using custom software. First, the electrocardiogram (EKG) signal was removed, the signal was bandpass filtered between 300-5000Hz, rectified, a 50ms median filter was applied, and the signal was downsampled to 1kHz. The amplitude of the diaphragm signal was then normalized by dividing by the standard deviation. Diaphragm bursts were detected using the scipy “find_peaks” function.

### Probe localization

Prior to inserting the probes into the caudal brainstem, they are dipped 5 times, for 5 seconds each dip, in DiI (Thermofisher #V22885), a red fluorescent dye, to mark the probe track. After the final recording of the procedure, the animal is euthanized, perfused with PBS, followed by 4% paraformaldehyde (PFA), and the brain is removed. The brain is fixed overnight in 4% PFA, 15% sucrose for 24 hours, and finally 30% sucrose. The brain is embedded in embedding medium (NEG-50) and frozen at −80C°. 25*μ*m coronal sections are collected, and imaged on an Olympus BX61VS slide scanner at 4x magnification. We then use SHARP-Track^26^ to downsample and manually transform each image to match a corresponding section of the Allen CCF. The red fluorescent probe tracks were identified in the registered slices and reconstructed in 3D atlas space. Special care was taken to accurately identify the probe tip position (i.e., the most rostral slice that DiI staining could be observed) to accurately reconstruct the rostro-caudal positioning of the probe, and the respective channels. We then visualized the activity of the probe alongside the extracted anatomical location to fine tune the rostrocaudal position of the probe.

### Anatomical subdivision of VRC and medulla

Since the preBötzinger Complex and Bötzinger Complex are not defined in the AllenCCF, we create a manual subdivision of the Paragigantocellular Reticular Nucleus, lateral part (PGRNl) to create these regions in our analyses. We define BötC as extending from the caudal edge of the facial nucleus (VII) (−5.96mm AP) for about 400*μ*m in the caudal direction (−6.4mm AP); preBötC as extending from this caudal edge (−6.4mm AP) to include the remaining PGRNl (−6.85mm AP). We restrict both preBötC and BötC such that the medial edge is in line with VII (0.89mm lateral). Any PGRNl units that do not fall into these defined regions remained labeled as PGRNl. Lastly, we combine the Superior Olivary Complex lateral (SOCl) and medial(SOCm) parts into one region (SOC).

### Coherence calculations

Coherence estimates whether two signals (here the breathing activity and a given neuron’s spiking activity) share frequency components. We use Chronux^44^ to compute the multi-taper coherence of the continuous integrated diaphragm signal (subsampled to 1kHz) with the point process signal of the spike times for a given unit within the first 5 minutes of a recording (coherencysegcpt). We use a time bandwidth product of 3 with 5 tapers. We compute the lower and upper bound on coherence using Jackknife error method with error bounds of [0.001, 0.999]. Chronux computes the phase lag of the unit (*ϕ*) at each frequency component. We determine the phase lag of a unit to be the value of *ϕ* at the frequency that is maximally coherent between the diaphragmatic signal and the unit’s spiking activity.

In addition to computing coherence, we also compute the directional selectivity index *L* ^73^ of each unit by:

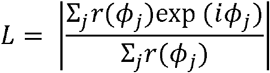

where *r*(*ϕ*) is the phasic firing rate of the unit as a function of respiratory phase, and *ϕ_j_* is the value of phase in the *j^th^* phase bin. *L* is 1-*σ*^2^ where *σ*^2^ is the circular variance of the firing rate as a function of respiratory phase.

### Phase computation and phasic responses

Phase (*ϕ*) is defined here from −*π* to *π* to normalize breaths across time and animals. We set the onset of each breath to *ϕ* = 0 and offset of each breath to *ϕ* = *π*. We then linearly interpolate a value of *ϕ* for each time sample between onset and offset. We set the sample immediately following breath offset to *ϕ* = −*π*, and linearly interpolate the samples until the next breath onset on the interval [–*π*,0). Phasic responses of individual neurons is computed as: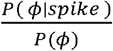. Note, the phase lag of a unit (defined in “Coherence calculations” is not necessarily the phase of the respiratory cycle for which that unit’s spike rate is maximal.

### Functional cell class decomposition

We perform unsupervised clustering to identify classes of neurons based on their phasic activity patterns. We first compute the phasic activity of a given unit as above (Phase computation and phasic responses) to get firing rate as a function of breathing phase. We bin the respiratory phase into 100 bins (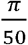 *radian* bin width). We then smooth the phasic curve with a 3 sample Savitzky-Golay filter, and divide by that units maximal firing rate. To account for possible dimensionality issues when clustering the phasic data, we create three sets of data to be clustered independently. The first set is the raw firing patterns which is a matrix **R** of size (*n* by *t*) where *n* is the number of units and *t* is the number of phase bins (100). We create the second set by performing PCA decomposition on the matrix **R**. We retaining only the leading four PCs, which account for >95% of the variance of **R** to create **P** which is of size (*n* by 4). We create the third set **N** by performing NMF decomposition on **R**. We retain 20 NMF components, such that **N** is of size (*n* by 20). We then cluster each set of data using KMeans clustering and Ward agglomerative clustering (*sklearn*). We vary the desired number of clusters from 2 to 15. We then compute the silhouette score for each combination of data preconditioning, clustering method, and number of desired clusters. Lastly, we repeat this analysis for subsets of **R** that include only units that have a lower bound of coherence value greater than a given threshold *T* where *T* = [0,0.1,0.3,0.5,0.7,0.9,0.99].

### Low dimensional population analyses

To compute the PCA decompositions of the populations, we first convert the spike times into a normalized, smooth spike rate by binning the spike rates into 5ms bins, smooth the bins with a Gaussian kernel *σ* = 10*ms* with and take the square root transform. The smoothed spike rates forms a matrix **X** of size (# neurons by # time bins) which is passed to the PCA. We fit PCA on data obtained during 100% O_2_ presentation, and apply that PCA decomposition to the entire recording.

For Extended Data Fig. 17 we compute the PCA decompositions for a given recording separately on each condition, which results in two square matrices **A, B** each of size (*n* by *n*) where n is the number of units. We compute the principal angles by first computing the singular value decomposition of the product of these two matrices:

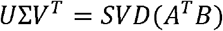

Where Σ is a diagonal matrix in which each the diagonal entries are the cosines of the principal angles *θ* between the subspaces, ranked by decreasing magnitude. We report the cos (*θ*) as “similarity”.

In addition to PCA, we also fit recurrent switching linear dynamical systems model (rSLDS)^48^ See: https://github.com/lindermanlab/ssm for detailed examples and explanations. Briefly, these models consist of *K* linear dynamical systems of the form:

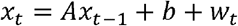

where *x_t_* is the low-dimensional continuous state vector of the system at time *t*, *A* is the matrix that specifies the continuous state update at each time step, *b* is a bias vector, and *W* is a noise term. *x_t_* and *b* have dimension *D*, and *A* is a [*D* × *D*] matrix. In the rSLDS, there are *K* number of *A* matrices and *b* vectors, one for each discrete state *k*, which we refer to as *A_k_* and *b_k_*. When the system is in discrete state *k*, the continuous state is now updated as:

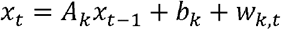

In comparison with a switching linear dynamical system (SLDS) in which the transition between the discrete states (*k*) is only dependent on the previous discrete state, switches between states in the rSLDS are dependent on the value of the continuous state *x_t_*. In other words, , the low-dimensional representation of neural activity will affect the probability of being in a discrete state. More precisely, the probability of the current discrete state *z_t_*, transitioning from discrete state *j* to discrete state *i* is given by:

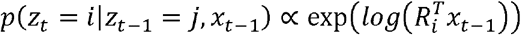

*R_i_* is a vector that weights the previous state *x*_*t*-1_.

We fit the rSLDS models with the number of discrete states *K* = 2 and the dimensionality of the continuous state *D* = 2. We use an isotropic Gaussian noise model for the dynamics. We employ a Poission emissions model so that the mapping between the continuous state x and the estimated neural activity *y_t_* (where *y_t_* is a vector of spike counts of length # neurons at time bin *t*) is a Generalized Linear Model with a Poisson distribution. We pass the spike counts in 10ms bins for all neurons during 100% O_2_ presentation as the inputs to fit the rSLDS model.

This fits two matrices *A* and bias vectors *b*—one for each of the *K* states—as well as defines the transition dynamics between these two dynamical systems. *A* and *b* define the flow fields in Fig. 2. We then supply an initial 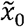 to generate simulated continuous states forward in time 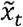 that depend only on the fit dynamics. We set the noise term *W* to zero after the observation that including a *W* term resulted in simulated states that were far more variable than the observed states, and that states simulated with omission of the *W* resembled the observed states. Lastly, we perform support vector regression (scikit-learn) to generate a mapping from the observed states *x_t_* to the observed integrated diaphragm activity. We then apply that mapping to the simulated continuous states 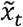 to generate a simulated diaphragm activity. The generative models either evoked a simulated diaphragm activity that exhibited periodicity and resembled normal diaphragm activity (Extended Data Fig. 12, regenerative), or did not generate periodic activity (non-regenerative). This distinction was clear, and no qualitatively in-between types of simulated diaphragm activity was observed. We tested whether a piecewise dynamical system with *K* = 2 discrete states was necessary and sufficient to result in a generative model capable of simulating diaphragm activity by fitting models with *K* = 1 and *K* = 3 respectively (Extended Data Fig. 12).

To align the latent dynamics of the rSLDS models, we first compute the average of the 2-d continuous state *x_t_* as a function of phase by computing 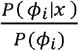 where *ϕ* is subdivided into 100 bins of size 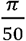 radians. This gives us a (100 by 2) matrix for each recording. We choose one recording to be the “target” recording, which we denote as *Z* and all others as source recordings, which we denote as *X*. We then compute the 2 by 2 alignment matrix *W* as:

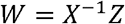

We then compute the aligned state: 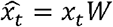

### Sigh and gasp detection

Sighs were detected by first computing the area under the curve (AUC) of the integrated diaphragm signal for each breath. We then compute the rolling median absolute deviance (MAD) with a centered 51 breath window. This gave, for each breath, the MAD of the 25 previous and 25 following breaths. Breaths for which the AUC of the integrated diaphragm was greater than 7 times the surrounding MAD were identified as sighs.

Periods of gasping were detected only during hypoxia presentation by first computing the interburst interval (IBI) for each breath. We smooth the IBI with a centered median filter with window length = 7 breaths. If the smoothed IBI exceeded 1s, a gasping period is determined to begin. The gasping period was deemed over if the smoothed IBI became shorter than 0.85s.

