## Extended Data Figures for "Latent neural population dynamics underlying normal breathing, opioid induced respiratory depression, and gasping"

### Extended Data Fig. 1

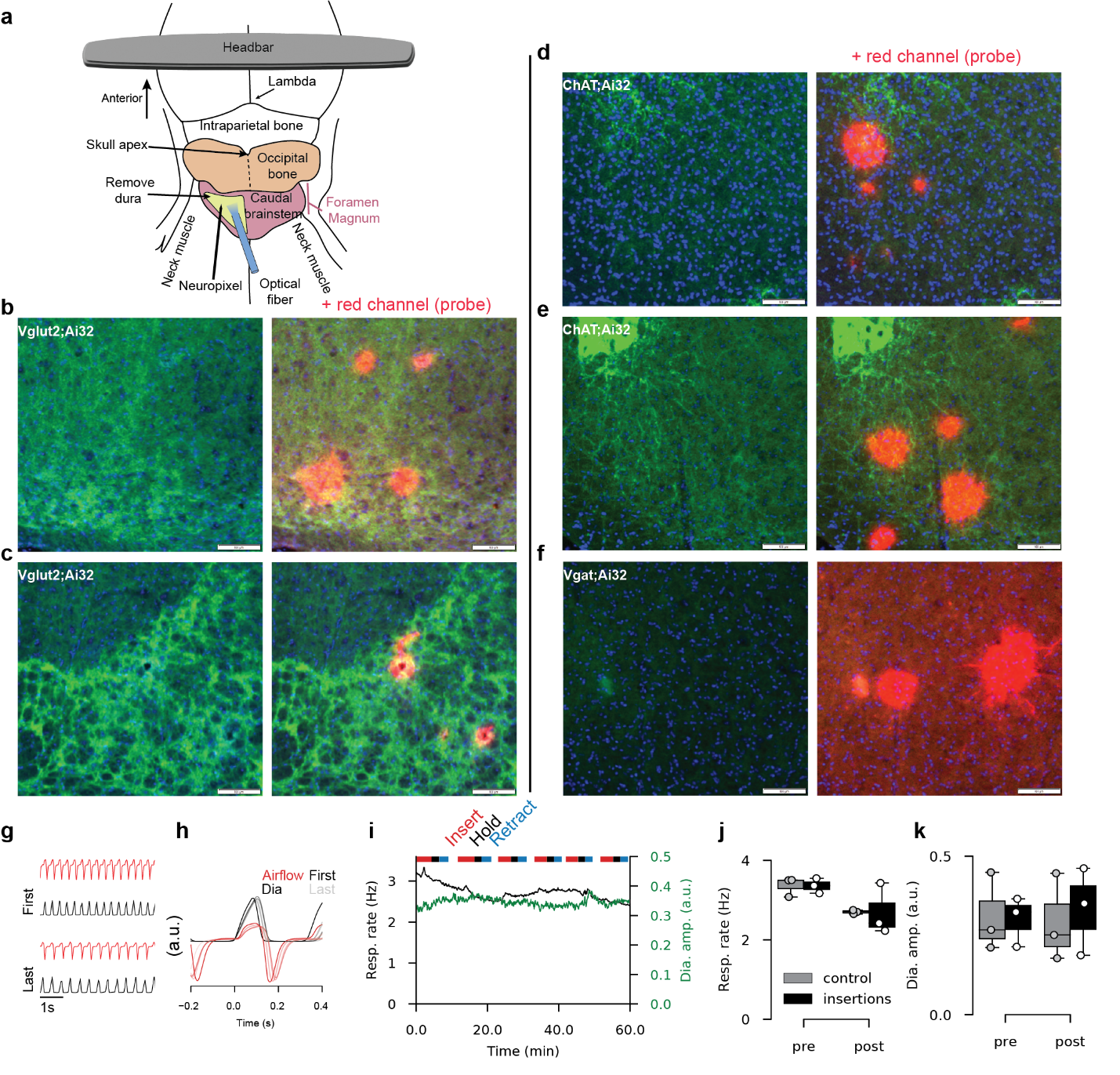

Surgical preparation causes minimal damage to VRC and alterations to normal respiratory behavior (**a**) Schematic of surgical preparation (See methods for details). (**b-f**) Five example coronal histological sections without (left) and with (right) the red channel visible. Blue channel is DAPI, green channel shows EYFP staining of transgenic expression of channel rhodopsin, red channel shows DiI tract indicating Neuropixel probe penetration. Probe penetrated in the anterior-posterior direction (perpendicular to the image plane), so each sequential coronal section shows an approximately circular diffusion of DiI. Smaller radii tracts indicate penetrations later in the experiment in which DiI has dissipated. (**g-k**) We perform separate test of the effect of repeated penetrations of the VRC on respiratory behavior. In 3 urethane anesthetized mice, we perform 6 sequential insertions of the VRC spaced in a 100-150$\mu m$ grid, centered around the VRC target as described in methods. Probes were inserted over approximately 5 minutes, held stationary for 3 minutes, and retracted over approximately 3 minutes. We monitored nasal airflow and diaphragmatic activity throughout. In 3 control mice we perform the same surgical and recording preparation including durotomy but perform no insertions of the Neuropixel probe. (**g**) Example airflow (red) and integrated diaphragm (black) after the first (top) and last (bottom) insertion of the probe. (**h**) Breath aligned average airflow (red) and integrated diaphragm (dia., black) after each sequential penetration. Later recordings are shown in increasing transparency. No qualitative differences are observed in respiratory behavior. (**i**) Respiratory rate (black) and integrated diaphragm amplitude (green) over the six sequential insertions in an example mouse (total time elapsed for each mouse was: [60,82,88] minutes). Horizontal lines at top of trace indicate insertion (red), holding (black), and retraction (blue) periods. A reduction in respiratory rate is observed, but no change in diaphragmatic amplitude is observed. (**j**) Respiratory rate and (**k**) diaphragmatic amplitude before (pre) and after (post) all insertions (black boxes: mean+/- IQR; white dots are individual mice). Mice in which no insertions were performed are compared at 90 minutes after initiation of recording (gray boxes: mean +/-IQR; gray dots are individual mice). Overall, insertions do not qualitatively alter breathing behavior. There are small transient increases in respiratory rate during insertion periods. Respiratory rate depression in mice without VRC insertions exhibit similar respiratory depression over time, likely due to prolonged urethane anesthesia. Thus, the insertion of the Neuropixel probe likely does not cause significant disruption of the VRC respiratory networks.

### Extended Data Fig. 2

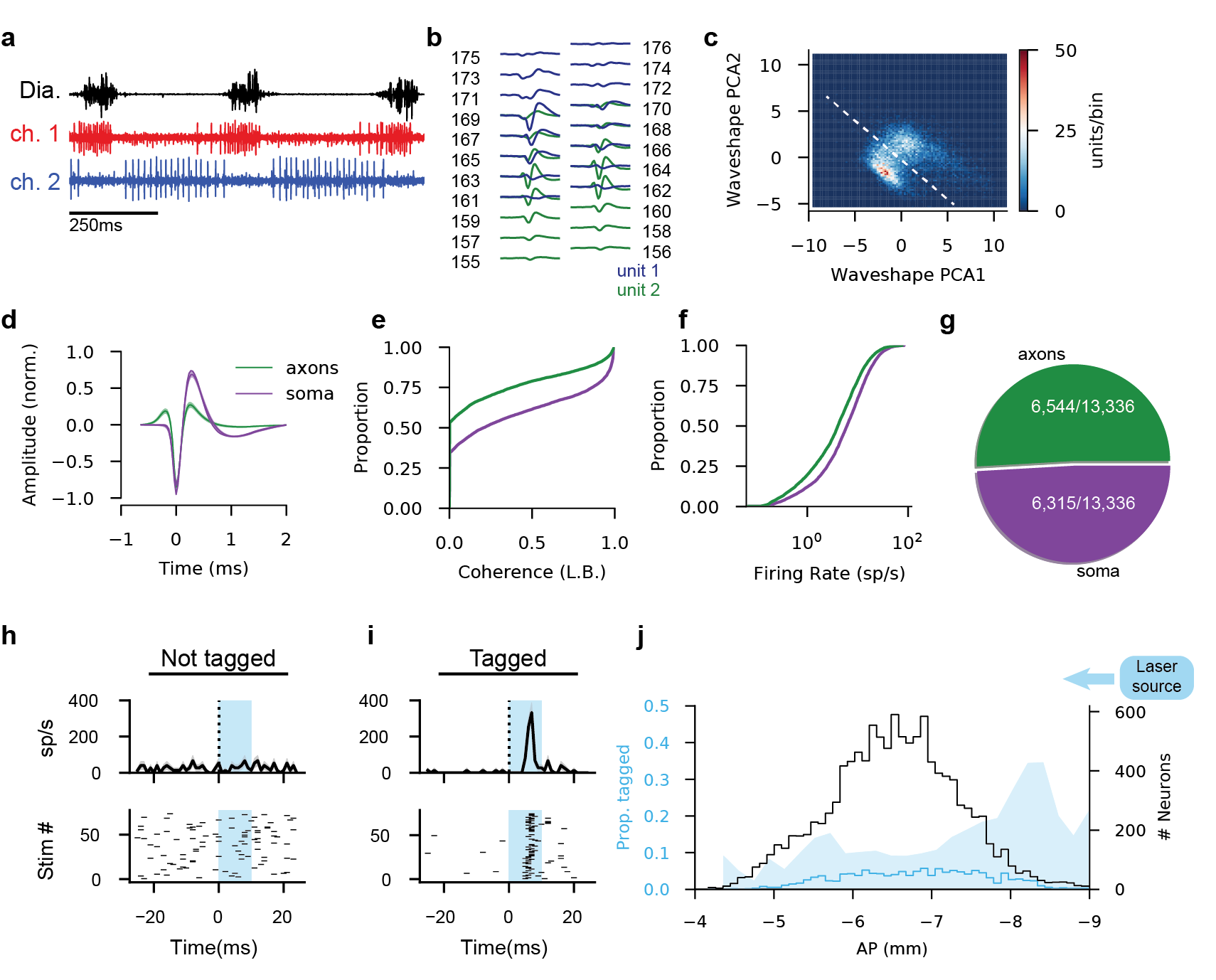

(**a**) Example raw signal from the diaphragm and two example channels with clear inspiratory and expiratory (red and blue, respectively) single units. (**b**) Average waveforms of two separable single units recorded simultaneously on overlapping electrode channels. Channel number is indicated. (**c**) PCA projections of waveshape features as detailed in ^1^. White dashed line is decision boundary separating putative axonal units from putative somatic units (**d**) Average waveform shape of putative axons (green) and soma (purple). Shaded regions are mean +/- 10*S.E.M. (**e**) Cumulative proportion of lower bound on coherence and (**e**) firing rate of axonal vs. somatic units. Somatic units are more likely to be strongly respiratory and have higher firing rates, suggesting the separation of axonal and somatic units is not arbitrary. (**g**) Proportion of axonal vs. somatic units of all single units obtained. (**h,i**) Optogenetic tagging results for an untagged neuron (h) and a tagged neuron (i). Shaded regions are presentation of a 10ms 473nm laser pulse with a sigmoidal on- and off ramp. Top: Stimulus aligned average firing rate ±S.E.M. Bottom: Stimulus aligned spiking for each of 75 laser pulses, each row is a stimulation, each bar is a spike. (**j**) Anterior-posterior distribution of positively tagged and untagged cells. Shaded region is proportion tagged (left abscissa), lines are absolute number of neurons (right abscissa) across all animals and genotypes. Laser is positioned posteriorly (right of histogram).

### Extended Data Fig. 3

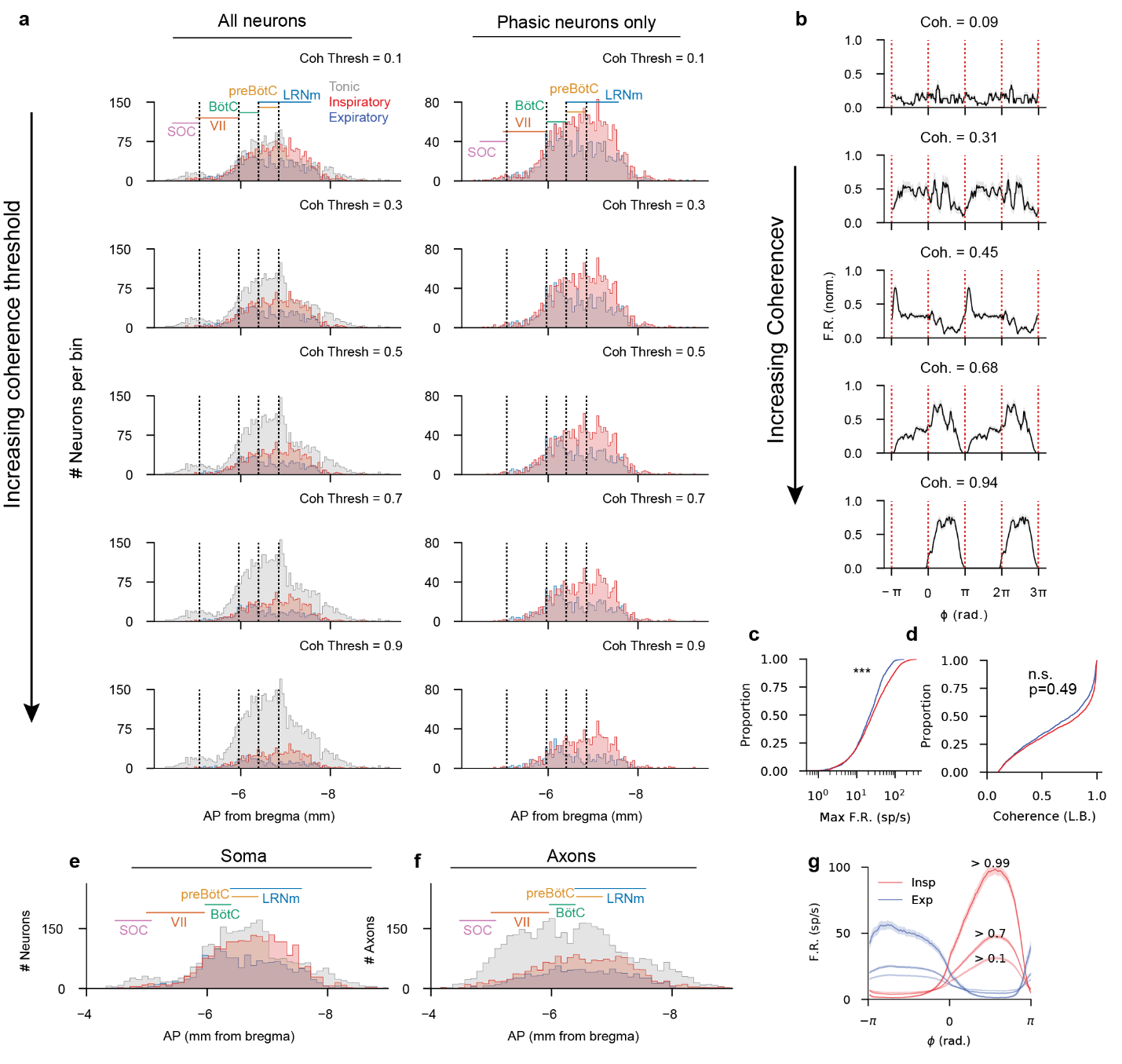
 Anatomical distributions are robust to coherence thresholds and the potential of contamination from non-local units. (**a**) Detailed anterior-posterior (AP) distributions of recorded units for varying coherence thresholds. (Left) All recorded units are classified as tonic (grey) inspiratory (red) or expiratory (blue) for increasing coherence thresholds (0.1, 0.3, 0.5, 0.7, 0.9). Units with a lower bound of coherence (C_lb_) less than the threshold value (i.e. weakly coherent units) are classified as tonic. Phasic units (i.e., units with a C_lb_ > threshold) were further classified as expiratory or inspiratory based on the phase lag of that unit relative to respiration. (Right) same as (left) but tonic neurons are omitted. Bins are 50μm. AP extent of named nuclei in the VRC are shown in the top row. There is a smooth anterior-posterior gradient in which expiratory neurons are found anteriorly and inspiratory neurons posteriorly. (**b**) Example phase aligned single unit firing rates at varying coherence values. Firing rates are normalized between 0 (no firing) and 1 (maximal firing rate) before taking the mean +/- S.E.M. Two cycles are shown. Inspiration onset (diaphragm activation) is $\phi=0,2\pi$; inspiration offset/expiration onset (diaphragm cessation) is $\phi=\pi,3\pi$. (**c**) Maximum firing rate and (**d**) C_lb_ cumulative distributions for all inspiratory (red) expiratory (blue) somatic units. (***p<0.001 Mann-Whitney U test) Inspiratory units have higher maximum firing rates than expiratory units. (**e**) AP distributions of somatic and (**f**) axonal units, tonic units are defined with C_lb_ threshold <0.1. Binsize is 150$\mu m$. Large densities of tonic axons are present rostral near the facial nucleus, and the increased density of expiratory units in the Botzinger region is not seen in axonal units. (**g**) Phase-averaged firing rates of all inspiratory (red) and expiratory (blue) units at three coherence thresholds (0.1,0.7,0.99).

### Extended Data Fig. 4

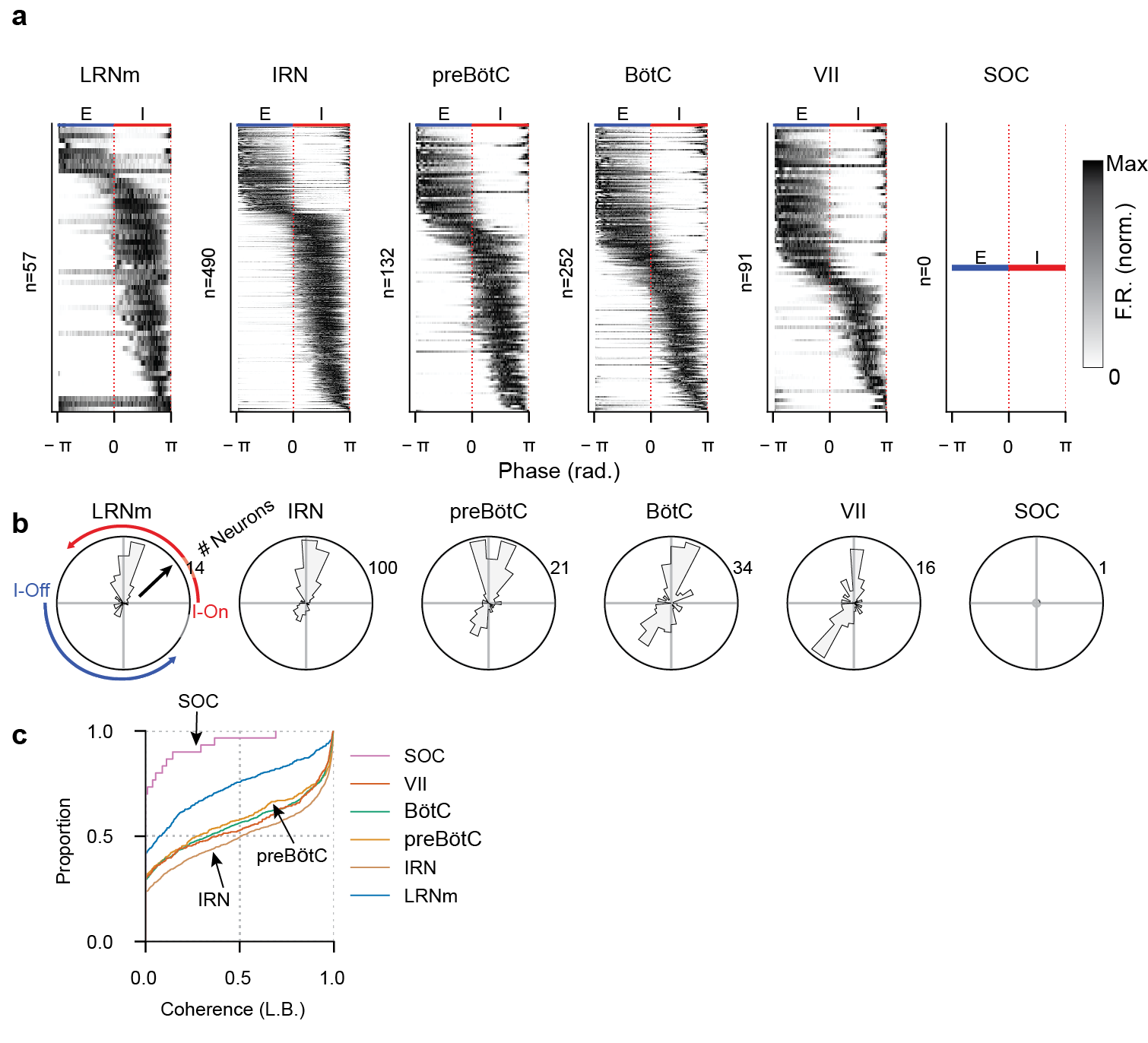

Phasic activity patterns of somatic units parcellated by VRC sub-regions. (**a**) Phase-averaged, maximum normalized activity of all strongly coherent (C_lb_>0.9) somatic units found in each VRC subregion. As expected, no SOC neurons are strongly coherent. Regions are arranged from caudal (left) to rostral (right). Proportionally more units are expiratory in more rostral regions. (**b**) Angular distribution of units with a given phase lag ($\phi$) with respect to breathing, for all regions in (a). Angular axis is respiratory phase, radial axis is number of units with a given $\phi$ value. Inspiration is top half of circle ($\phi>0$), expiration is bottom half of circle ($\phi<0$) (**c**) Cumulative distributions of the C_lb_ values for all somatic units in each region. Rightward shifted curves indicate a higher proportion of coherent units.

### Extended Data Fig. 5

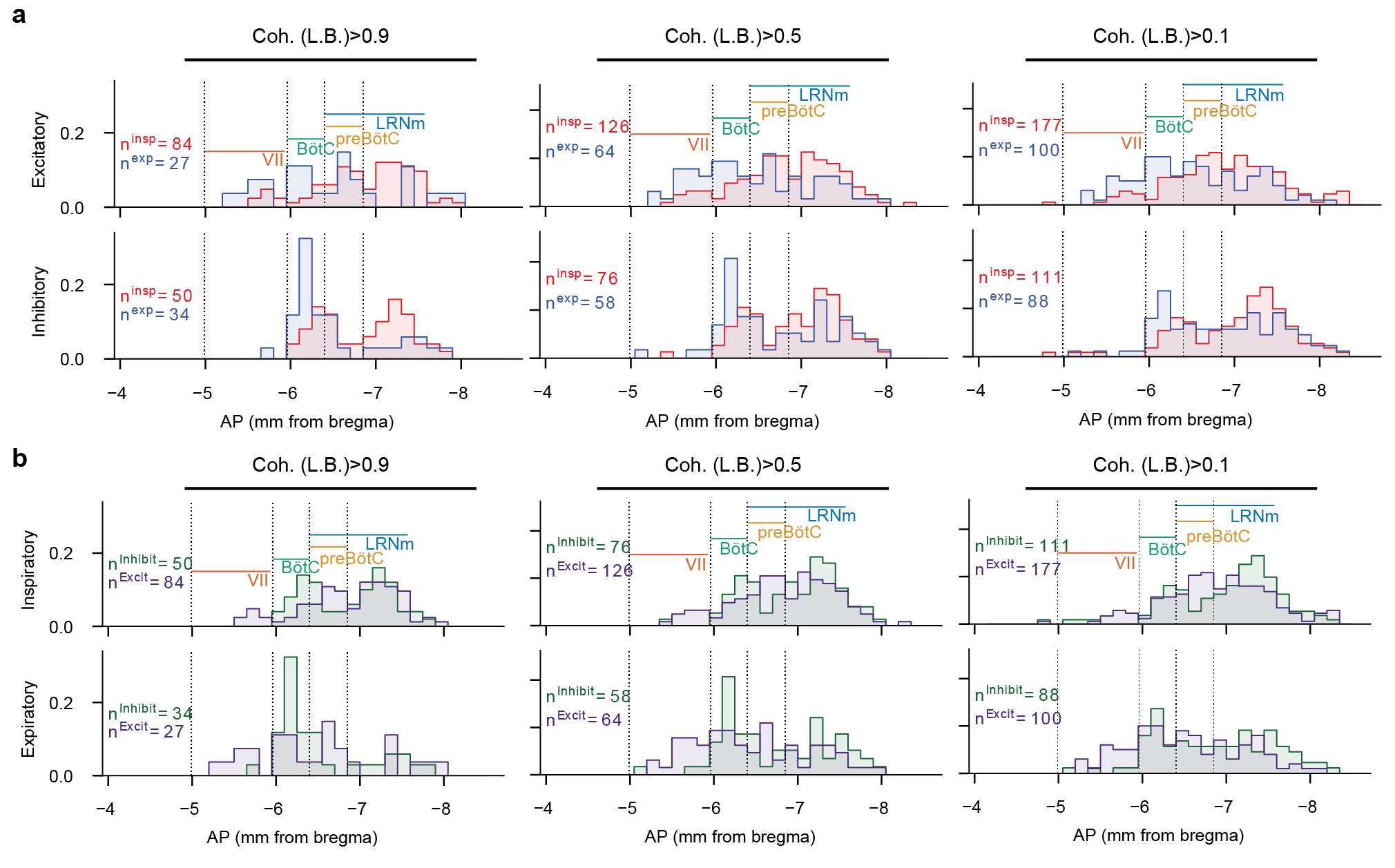

Anterior-posterior distributions of somatic units of identified genotype respiratory activity pattern. Given that Dbx1^+^ neurons are glutamatergic, we combine Dbx1^+^ opto-tagged neurons with Vglut2^+^ opto-tagged neurons to identify the excitatory units. (**a**) AP distributions of inspiratory (red) and expiratory (blue) units that are excitatory (top) or inhibitory (bottom) for varying coherence thresholds (left to right). Y-axis is probability of a unit being in a given bin (150$\mu m$), normalized within the identified group (i.e., inspiratory or expiratory are normalized independently). Of note is the high proportion of inhibitory neurons in the canonical Botzinger complex, particularly if analyses are restricted only to strongly coherent neurons, and the high proportion of inspiratory neurons caudal to the preBötC, likely including bulbospinal rVRG premotor populations. (**b**) Same data as in (**a**), but colored by excitatory (purple)/inhibitory (green); inspiratory neurons in top row, expiratory neurons in bottom.

### Extended Data Fig. 6

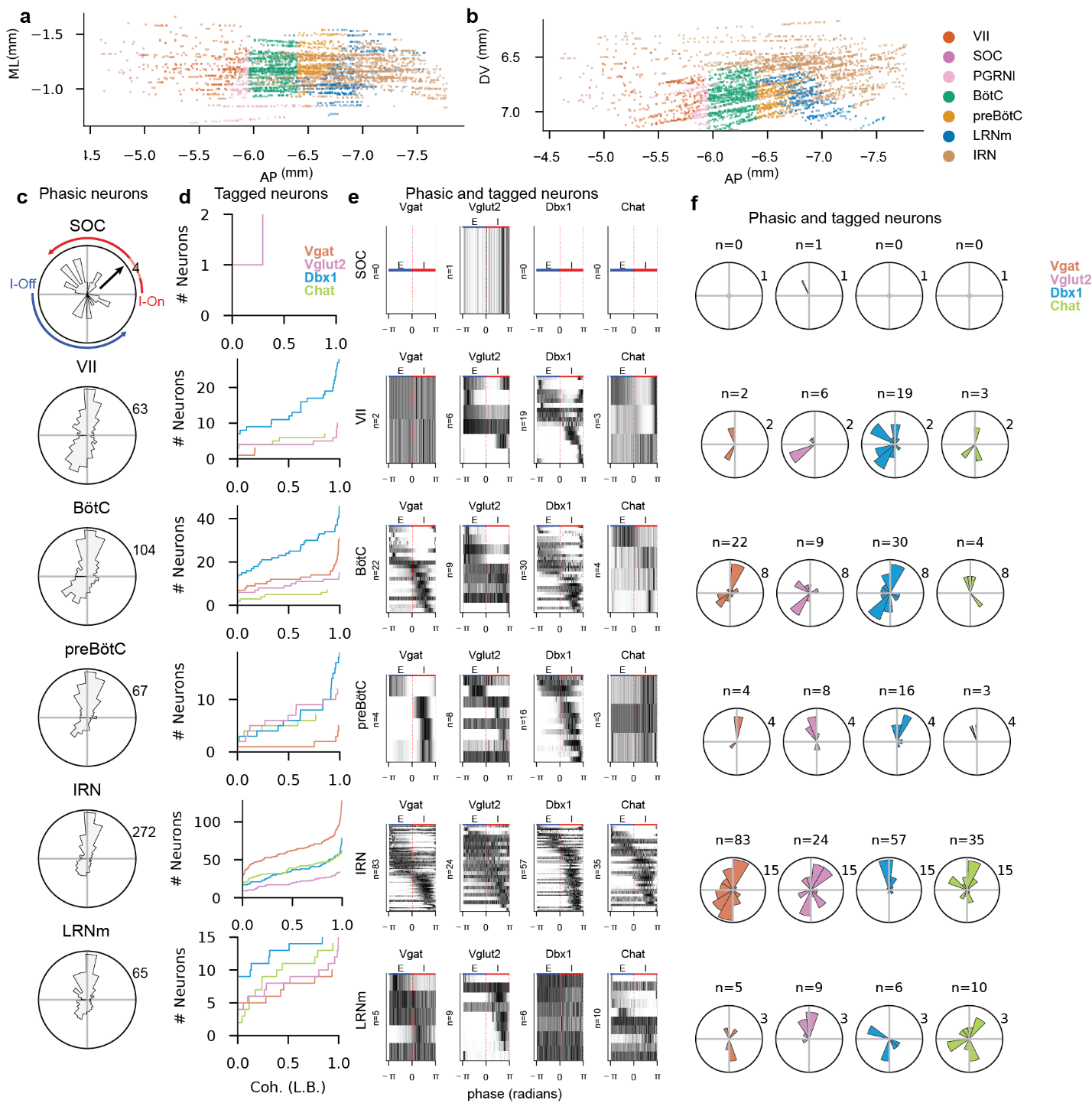

Detailed functional attributes of anatomically identified and optogenetically tagged units. (**a,b**) As the preBötC and BötC are not defined in the AllenCCF, we subdivide the Paragigantocellular Reticuclar Nucleus (lateral) (PGRNl) region into preBötC and BötC (See Methods) (**a**) Horizontal and (**b**) saggital projection of all recorded units colored by their identified subregion. (**c**) Distribution of preferred phase of all phasic units ($C_{lb}$>0.1) in the given region. Proportionally more preBötC neurons are inspiratory. Radial axis is number of neurons; angular axis is preferred phase. (**d**) Cumulative counts of lower bound of coherence values for all opto-tagged units in a given region, colored by their genotype. Y-axis is number of units. Right shifted curves represent a greater proportion of the population are strongly coherent (e.g., Vgat^+^ BötC units are strongly coherent as a population). (**e**) Phasic activity patterns of all phasic ($C_{lb}$>0.1) opto-tagged units in each region. Each panel is ordered by preferred phase. (**f**) Polar histograms of preferred phase for all phasic, opto-tagged units in (e); angular axis is preferred phase, radial axis is number of neurons as in (c).

### Extended Data Fig. 7

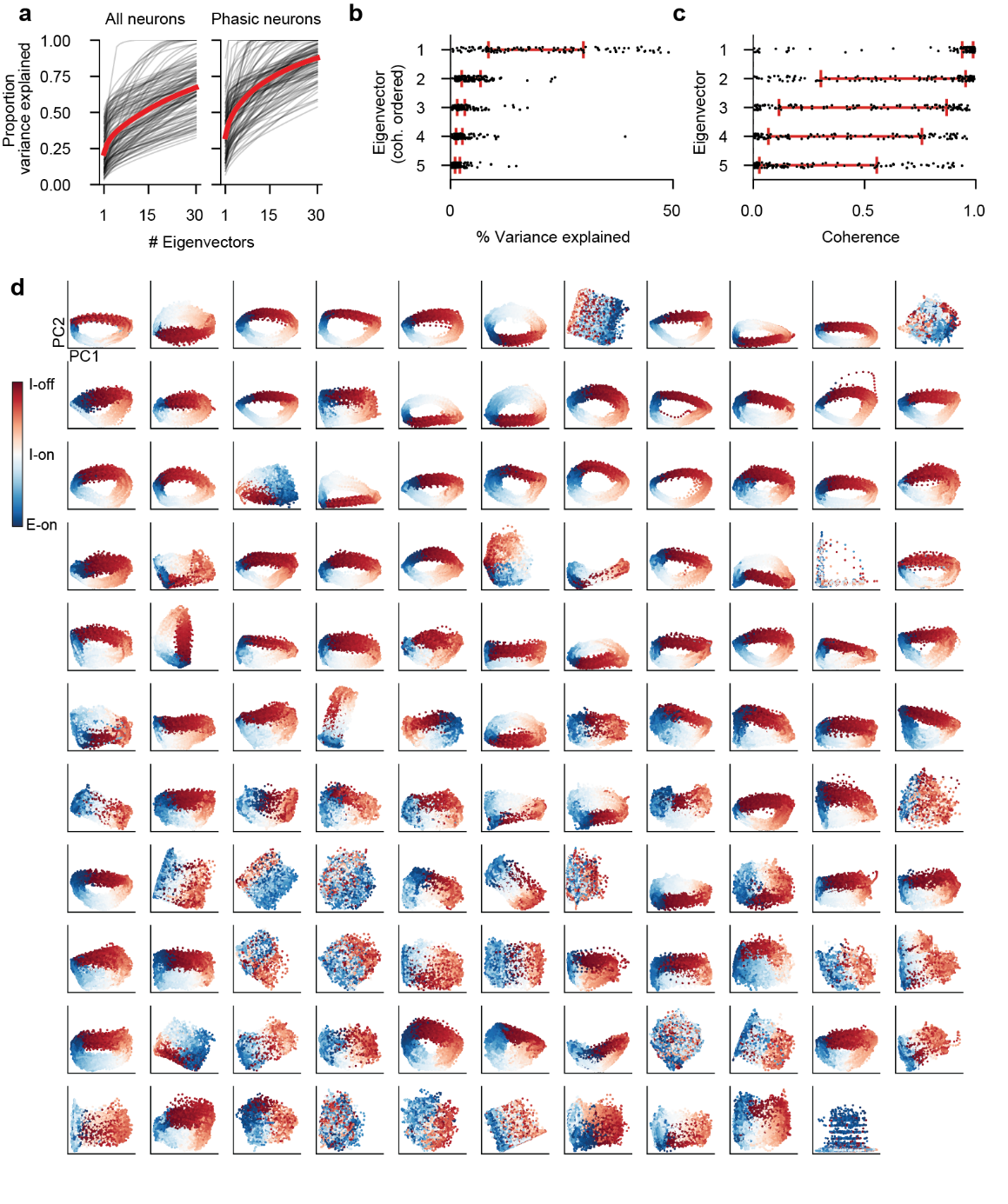

PCA decompositions of simultaneously recorded neural populations. (**a**) Cumulative explained variance as a function of number of PCs included. Black lines are individual recordings, red lines are mean across all recordings. Left is PCA decompositions which include all neurons, right is PCA decompositions that include only phasic neurons ($C_{lb}>0.1$). (**b**) We reorder the PCs not by their eigenvalue but by the coherence of the PC with breathing in order to compare across recordings. The percent variance explained for eigenvectors in descending order of coherence is shown. After re-ordering the PCs, on average the eigenvectors decrease in their explained variance ratio monotonically, suggesting that the coherence re-ordered PCs still capture a large amount of the variance of the neural population in the leading PCs. Black dots are individual recordings, red errorbars are median +/- IQR (**c**) Similar to (b) we show the coherence value for the leading five eignevectors (now ordered by eigenvalue). On average, the leading eigenvectors are most coherent with breathing. (**d**) Projections of population activity onto the two most coherent PCs for all recordings. Many recordings exhibit similar rotational structure that is highly consistent. Further, most recordings show a strong mapping between position on the manifold and respiratory phase (red->blue).

### Extended Data Fig. 8

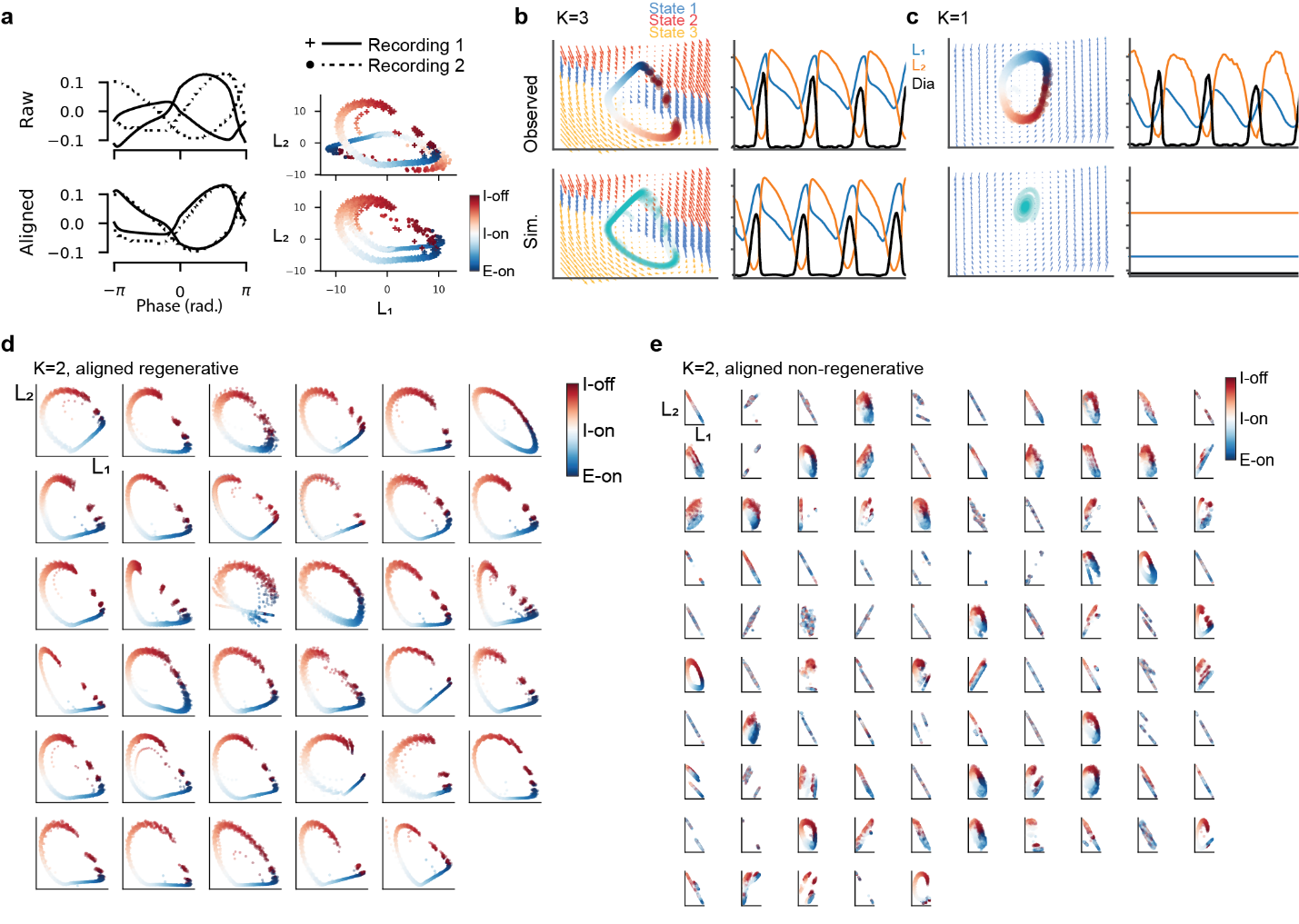

Recurrent linear dynamical systems uncover conserved latent dynamics that switch between two states if sufficient populations are recorded to recreate regenerative dynamics. (**a**) The latent structure of 2 separate recordings may have similar qualitative characteristics, but differ in the permutation and scaling of the latent axes (top). We linearly re-align the two recordings (see Methods) such that the trajectories overlap one another (bottom). Left shows the phase aligned average of the two latent components for two recordings before (top) and after (bottom) linear alignment. The phase-colored trajectories through the unaligned (top) and aligned (bottom) latent spaces. Dots are 5ms time bins. (**b**) Increasing the number of states (i.e., partitions of the latent space with different dynamics) for the rSLDS does not change the qualitative structure of the latent trajectories, and does not drastically increase the number of recordings that are regenerative (35 for $K=2$, 37 for $K=3$).(**c**) Allowing no partitions ($K=1$) of the latent space prevents any recording from being regenerative (example shown top), despite oscillatory structure in the observed data. Rather, single state LDSs form decaying spirals. Panel organized as in Fig. 4a-d. (**d**) The trajectories of all regenerative recordings, and (**e**) non-regenerative recordings, aligned to a reference recording. Markers are colored by phase as in (a)

### Extended Data Fig. 9

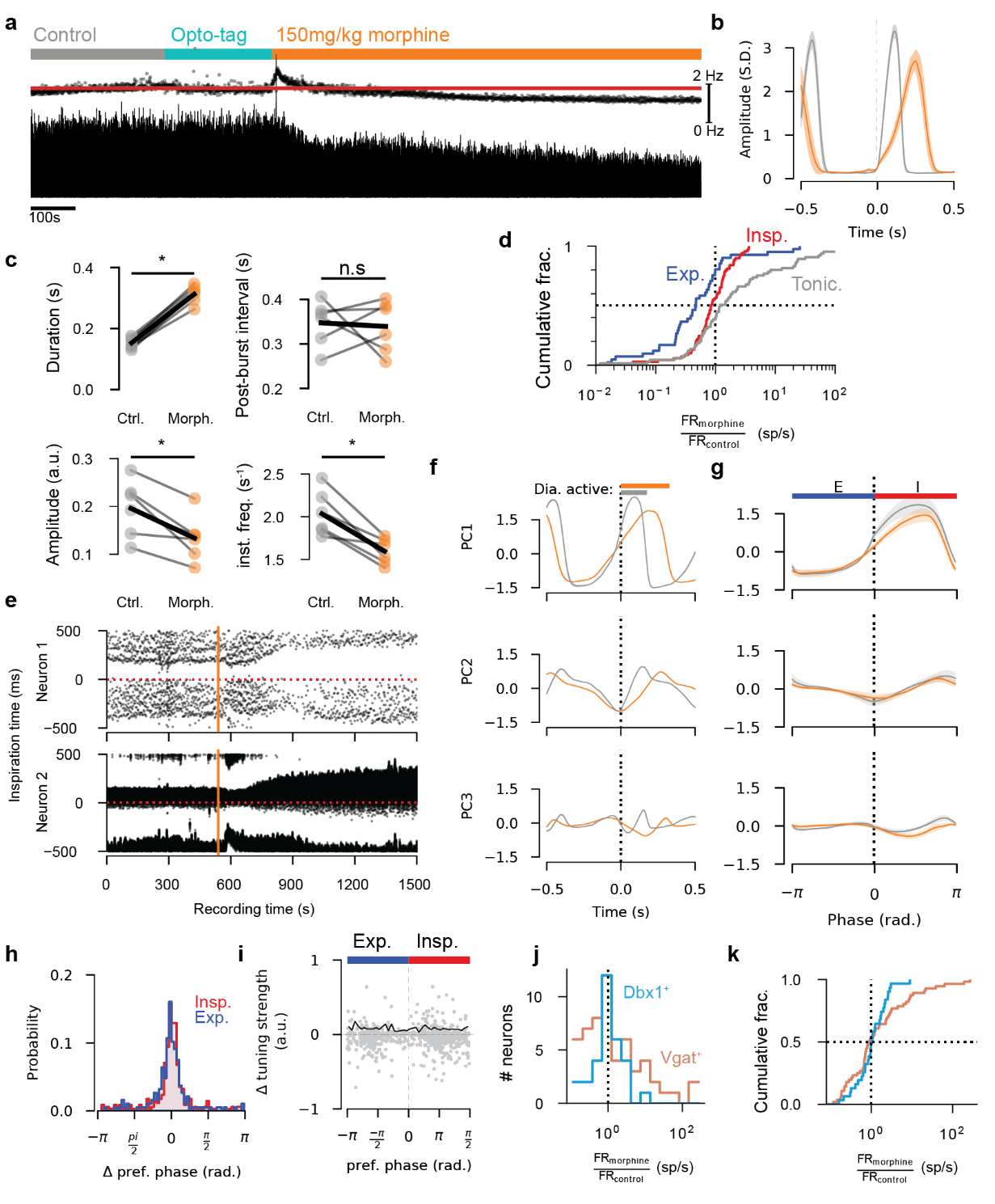

Morphine effects on respiratory and neural activity (**a**) Example integrated diaphragm trace before and after morphine administration. Top scatter is instantaneous respiratory rate. Red line is average respiratory rate in control period. Scale bar for respiratory rate is right (0-2Hz). Opto-tagging period interleaves control and morphine administration. (**b**) Average breath onset aligned diaphragm activity in control and morphine (taken 10 minutes after initial morphine administration). Shaded regions are mean $\pm$S.E.M. (**c**) Changes in diaphragm amplitude, burst duration, respiratory frequency, and interburst interval as a result of morphine administration for all recordings with morphine administration (n=6). Morphine slows respiratory rate by increasing diaphragm contraction duration. (*p<0.05 2-sided paired t-test) (**d**) Cumulative probability distribution of the ratio of firing rates in morphine to firing rates in control (FR_ratio_), for all recorded neurons. Expiratory neurons are de-recruited on average (**e**) Breath onset aligned rasters for an example expiratory (top) and inspiratory(bottom) over the course of morphine administration. X-axis is time in the recording of a given breath, y-axis is breath onset aligned spike time. Red dashed horizontal line is breath onset time. Each column is a breath and each dot is a spike. Morphine is administered at the vertical orange line. (**f**) Breath onset aligned average of the first 3 PCs of neural population in control and morphine for an example recording. Morphine dramatically slows the evolution of the population activity. (**g**) Phase-aligned average PCs of neural population averaged across all recordings. Shaded region is mean$\pm$S.E.M. Phasic structure of the PCs is preserved in morphine, with a slight reduction in amplitude of PC1. (**h**) Histogram of change in preferred phase of all recorded neurons. Preferred phase generally is stable for both inspiratory and expiratory neurons after morphine administration. (**i**) Change in respiratory tuning strength as a function of preferred phase. Each dot is a neuron, solid black line is the mean absolute value of the change in tuning strength. Neither expiratory nor inspiratory neurons tend to change their tuning strength. (**j**) Histogram and (**k**) cumulative distribution of FR_ratio_ for optogenetically identified neurons.

### Extended Data Fig. 10

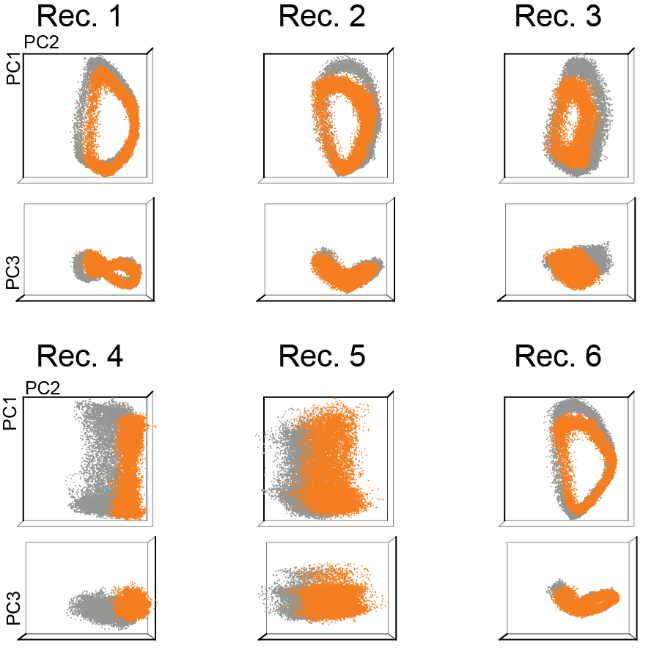

Projections of the population activity onto the leading three PCs for all recordings in which morphine was administered. Grey dots show control period, orange dots show after morphine administration. Recordings which show rotational activity are largely preserved in morphine, while those with little rotational activity show larger changes suggesting that morphine acts not to disrupt the oscillatory behavior of the population, but rather affects the non-oscillatory modes.

### Extended Data Fig. 11

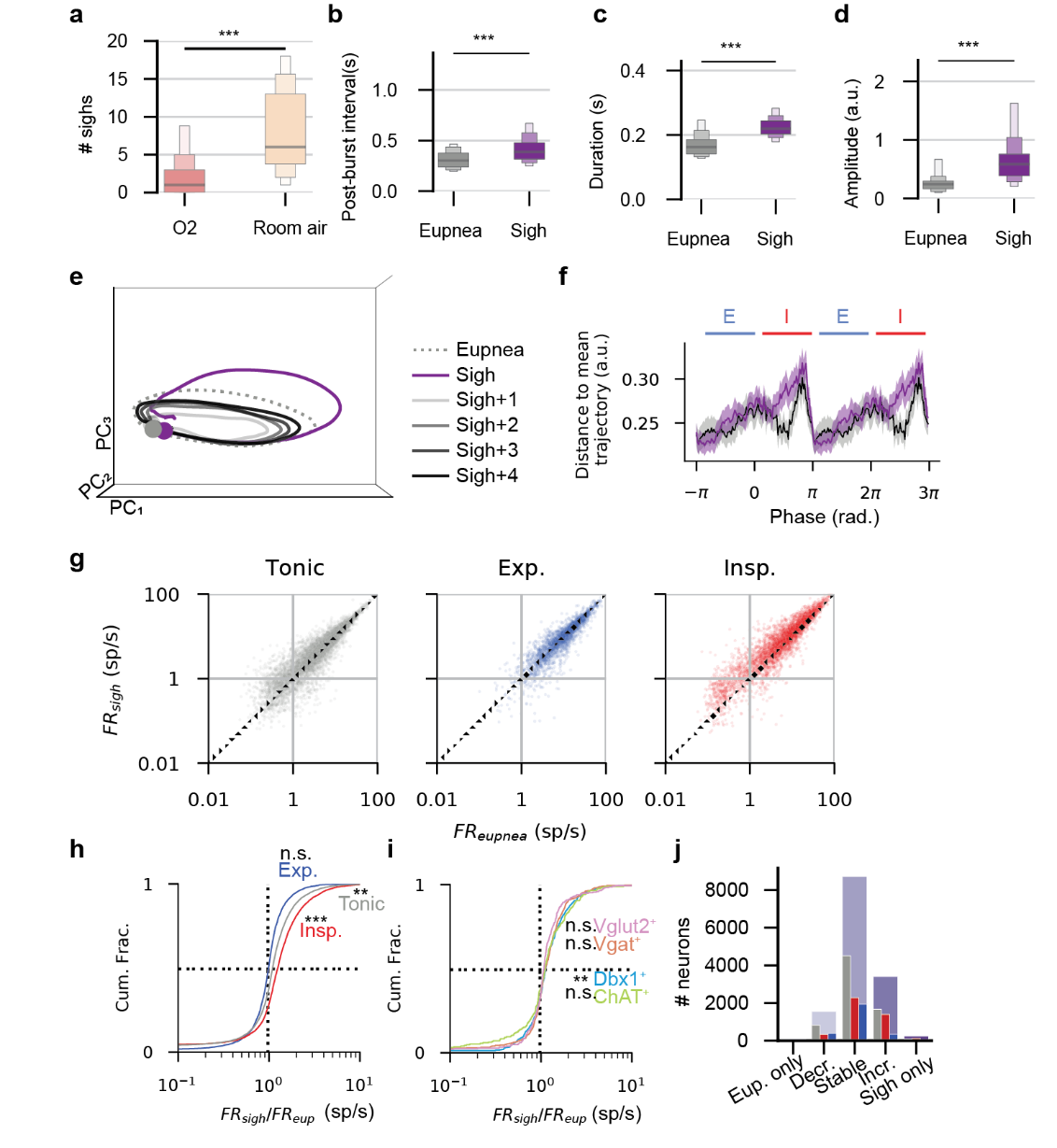

(**a**) Average number of sighs observed in 100% O_2_ and in room air across all recordings (paired t-test ***p<0.001). (**b**) Average post-burst interval (time from breath cessation to onset of next breath), (**c**) breath duration, and **(**d) breath amplitude are increased during sighs (paired t-test ***p<0.001). (**e**) Mean neural PC trajectories as in Fig. 6f, but including the first through fourth eupnea after the sigh. Recovery to a normal eupneic trajectory takes several breaths after a sigh. (**f**) Distance to mean eupnea trajectory as in Fig. 3i, averaged across all recordings and breaths. Shaded region is mean $\pm$S.E.M. Distance from sigh to eupnea shown in purple. Sighs follow the same trajectory as eupnea except during an excursion in the late inspiratory burst period. (**g**) Firing rates for all units across all recordings during eupnea (x-axis) and sighs (y- axis). Each dot is a unit, units are separated by their preferred phase in eupnea. Somatic and axonal units are shown (**h**) Cumulative probability distribution of the ratio of firing rates in sigh to firing rates in eupnea (FR_ratio_), for all recorded neurons. Inspiratory and tonic neurons are right shifted (i.e., recruited) during sighs (*** p<0.001, **p<0.01, 1-sample t-test on log firing rate ratio). (**i**) As in (h) for neurons of optogenetically identified as indicated by color. Dbx1^+^ units are weakly recruited during a sigh (p<0.01, 1 sample t-test on log firing rate ratio). (**j**) Neurons are categorized by the FR_ratio_ as being eupnea only (FR_ratio_<0.2), sigh decreasing (0.2$\leq$FR_ratio_$<$0.7), stable (0.7 $\leq$FR_ratio_$<$1.5), sigh increasing (1.5$\leq$FR_ratio_$<$5), or sigh only (5$\geq$FR_ratio_). The total number of neurons in each category are shown in purple, and are subdivided by inspiratory, expiratory and tonic. Few neurons are eupnea or sigh only, while most are relatively stable. ***p<0.001 2 sided t-test for panels a-d. Boxen plots are median, boxes are median $\pm$first, second, and third quantile respectively.

### Extended Data Fig. 12

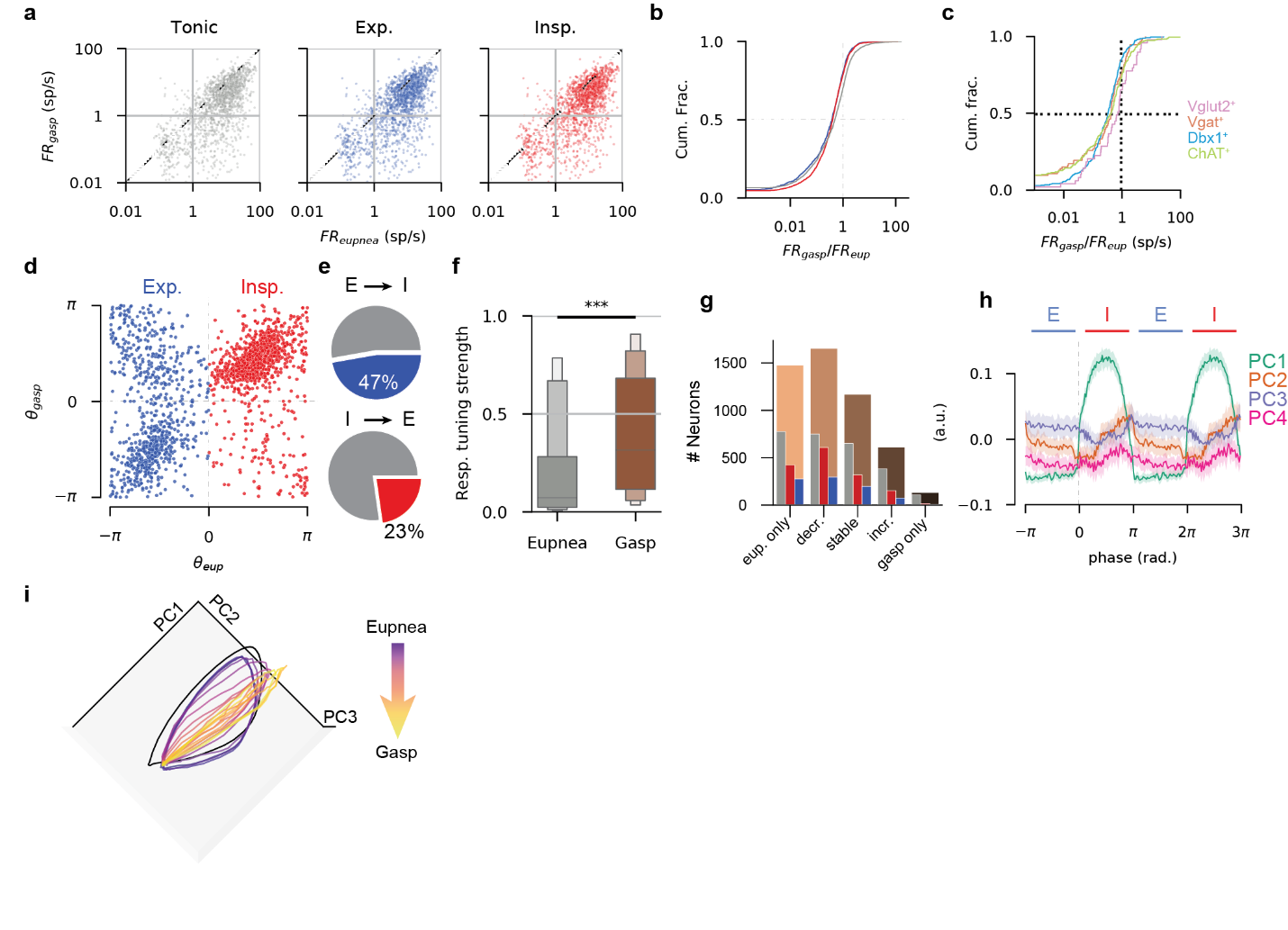

(**a**) Firing rates for all units across all recordings during eupnea (x-axis) and gasps (y- axis). Each dot is a unit, units are separated by their preferred phase in eupnea. Somatic and axonal units are shown (**b**) Cumulative probability distribution of the ratio of firing rates in sigh to firing rates in eupnea (FR_ratio_), for all recorded neurons. All populations are left-shifted (i.e., reduced firing rates) during gasping. (p<0.001 1-sample t-test on log firing rate ratio). (**c**) As in (b) for neurons of optogenetically identified genotype as indicated by color. An optogenetically identified unit’s identity is not indicative of whether it will be recruited or de-recruited during a gasp. (**d**) The preferred phase of each neuron ($\theta$) is plotted as computed during eupnea (x-axis) and gasping (y-axis). Neurons along the diagonal show stable phase preference, while neurons off the diagonal show switching of preferred phase. (**e**) 47% of expiratory units became inspiratory during the gasp, while 23% of inspiratory units became expiratory. (**f**) Distribution of respiratory tuning strength of all units during eupnea and gasping. On average units become more tuned to respiration during gasping. Boxes are median $\pm$first, second and third quantile, respectively. (p<0.001 Wilcoxon rank-sum test) (**g**) Neurons are categorized by the FR_ratio_ as being eupnea only (FR_ratio_<0.2), gasp decreasing (0.2$\leq$FR_ratio_$<$0.7), stable (0.7 $\leq$FR_ratio_$<$1.5), gasp increasing (1.5$\leq$FR_ratio_$<$5), or gasp only (5$\geq$FR_ratio_). The total number of units in each category are shown in brown, and are subdivided by inspiratory, expiratory and tonic. In general, neurons are de-recruited during the gasp. (**h**) Breath-aligned average first 3 principal components for an example recording during eupnea and gasping. Pre- and post- inspiratory population activity is largely lost during gasping. (**i**) The collapse from rotational to ballistic dynamics is gradual. Each line is the average trajectory in sequential 10s periods during the transition from eupnea (purple) to gasping (yellow). The black trajectory is the average eupneic trajectory prior to hypoxia exposure.

### Extended Data Fig. 13

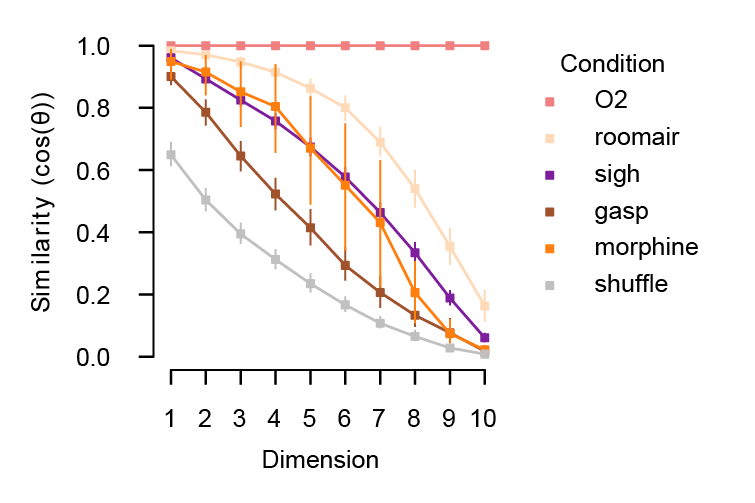

Low-dimensional structure is most disrupted during gasping. We fit PCA separately during different conditions and compute the principal angles between the subspaces spanned by the leading eigenvectors for each condition. Similarity (defined as the cosine of the principal angles) between the leading dimensions of the PCA decomposition for each condition as compared to control (100% O_2_). Unsurprisingly, the similarity is highest for the room air condition. Sighs and OIRD (morphine) are next similar, and gasps are least similar. To compute the lower bound on similarity, we shuffle the entries of the eigenvectors for O_2_ and compute the principal angles between the shuffled space and the original space (gray). Markers are median and lines are +/- IQR.

### Extended Data Fig. 14

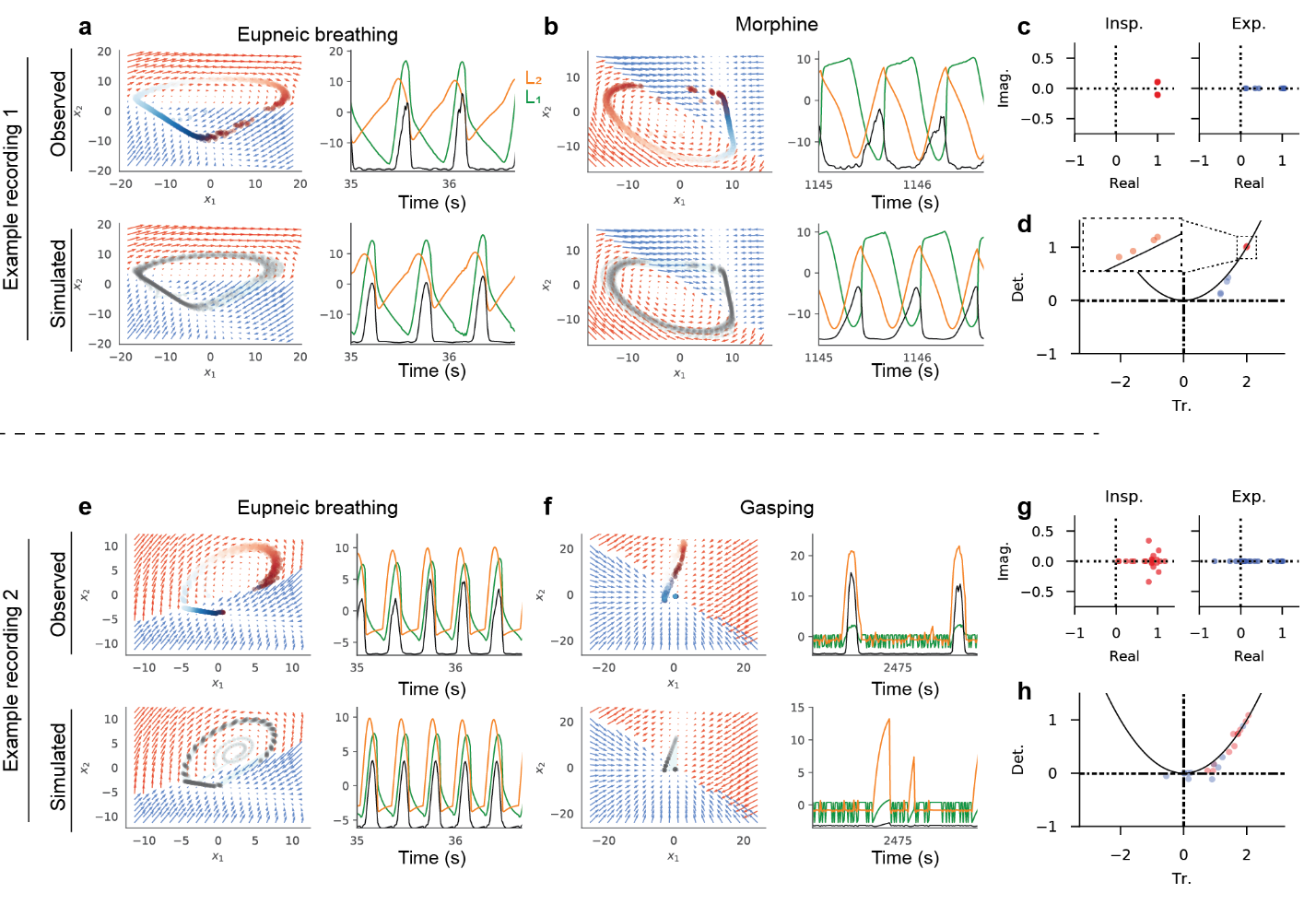

Maintenance of low-dimensional structure during morphine administration, and loss of that structure in gasping, is corroborated using rSLDS models. (**a**) rSLDS fit and simulation for an example regenerative recording of eupneic breathing. Layout as in Fig. 4e. (**b**) rSLDS fit and simulation for the recording in (a), during moprhine. Rotational, regenerative dynamics are maintained during morphine, although the axes of the latent space have been rotated. (**c**) Eigenvalues of the rSLDS dynamics matrices as fit during opioid administration. Only recordings that were regenerative in control are shown. Dots are colored by inspiratory (red) or expiratory (blue) state. Inspiration has imaginary eigenvalues, while expiration does not. (**d**) Same as Fig. 5l, recreated for context. Black parabola is $Tr^{2}=4\cdot Det$. (**e-h**) as in (a-d), for a different regenerative recording that was subject to hypoxia. Note the loss of rotational dynamics during gasping. (h) is same as 7h, recreated for context.

### Extended Data Table 1

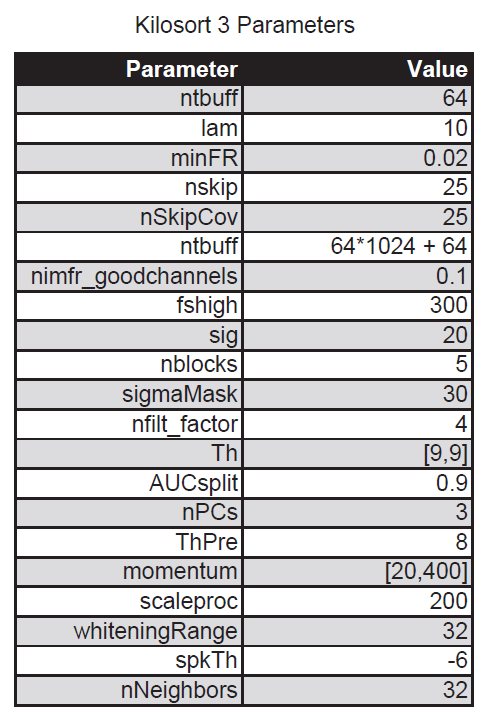

Kilosort 3 parameters

### Extended Data Table 2

| Abbreviation | Region |
| --- | --- |
| BötC | Bötzinger Complex |
| IRN | Intermediate Reticular Nucleus |
| LRNm | Lateral Reticular Nucleus, medial part |
| PGRNl | Paragingatocellular Reticular Nucleus, lateral part |
| preBötC | pre-Bötzinger Complex |
| SOC | Superior Olivary Complex |
| VII | Facial Nucleus |

Anatomical region abbreviations, modified from the AllenCCF.

### Extended Data Video 1.

Anatomical locations of all recorded neurons. The outline of the entire brain, as well as the facial nucleus (VII) and nucleus ambiguus are shown. Gray neurons on the left side are tonic neurons, light purple neurons on the right side are weakly phasic neurons (0.1 $<$ coherence lower bound $<$ 0.9), and dark purple neurons on the right side are strongly phasic neurons(coherence lower bound $>$ 0.9). Note that the strongly phasic neurons are found in a more restricted region of the ventrolateral medulla than either the tonic or weakly phasic neurons and are not often found within VII. Visualization performed with Brainrender^2^

### Extended Data Video 2.

Neural population activity evolves through constrained trajectories in a low-dimensional space. Integrated diaphragmatic activity (top) and (middle) activity of the 249 simultaneously recorded neurons in Fig 2. evolve over time. Firing rates are binned at 5ms and smoothed with a Gaussian kernel of s.d. = 15ms. Firing rates are normalized to maximum firing rate for visualization. Purple is 0 firing rate, yellow is maximal. Trajectories evolve over time through constrained PC space (bottom). Color of moving trace in bottom indicates diaphragmatic activity (black is inactive, yellow is maximally active). Video speeds from 15% real time to 50% real time after several seconds of playback. The inspiration off attractor described in Fig. 2 is appreciated in the temporal evolution of the neural population trajectories. A sigh and associated post-sigh eupnea is observed early in the video; the neural population trajectory exhibits an excursion from the typical eupnea trajectory.

### Extended Data Video 3.

Regions of a stable neural manifold correlates to multiple respiratory features. Each quadrant of the video shows the position of the neural population in the space defined by the leading 3 PCs for a 1000s time period, with each dot representing a single 5ms time bin. Quadrants are colored by the (top left) diaphragmatic activity, (top right) trajectory speed, (bottom left) 4^th^ PC, and (bottom right) respiratory phase.

### Extended Data Video 4.

Morphine slows neural population trajectories through PC space. Red line shows trajectory evolution at 30% real time speed in control (left) and morphine (right). Colored dots show 50s of overlapping neural states, where each dot is a 5ms bin. Dots are colored by trajectory speed.

### Extended Data Video 5.

Transition from eupnea to gasping and recovery. Video laid out as in Supplementary Video 1. 97 neurons are shown in the raster. Video plays at 100% real time of recording. Rotational trajectories through the PC space are observed during eupnea. These rotations become ballistic efforts during gasping (brown traces). During recovery, the trajectories (blue) smoothly transition back to normal eupnic rotations.
